# Stochastic foraging paths primarily drives within-species variations of prey consumption rates

**DOI:** 10.1101/2024.05.29.596370

**Authors:** Vincent Bansaye, Geoffroy Berthelot, Amina El Bachari, Jean-René Chazottes, Sylvain Billiard

## Abstract

The speed at which individuals interact, in particular prey and predators, affects ecological processes at all scales, including how fast matter and energy flow through ecosystems, and how stable communities are. Environmental heterogeneity and individual variabilities are generally believed to be the main factors underlying the variation of consumption rates of prey by predators. We challenge this view by comparing predicted variability from a stochastic model to experimental data. We first analyze a stochastic model of a simple random walk with elementary ecological processes involved in prey consumption, including prey depletion, predator movements and prey handling. We provide sharp approximations of the distribution of the consumption rate and a quantitative prediction of the coefficient of variation when stochastic foraging is the only source of variability. Predictions are then compared to the coefficients of variation estimated from data from dozens of various species and experimental contexts. We show that the predictions only accounting for intrinsic stochasticity in foraging are compatible with the range of observed values, in particular in 1 or 2 dimensional space. After evaluating the robustness of our model’s predictions through stochastic computer simulations, we conclude that the main driver of the variation of the consumption rate is the foraging process itself rather than environmental or between-individual variabilities. Our approach lays the foundations for unifying foraging theory and population ecology, and as such has many empirical and theoretical implications for both fields.

---

The assessment of ecosystems services, of the impact of harvesting natural resources, or of the stability of ecological communities requires identifying the main drivers of biomass and energy flux and species dynamics. In particular, the consumption rate of prey at different levels of trophic food webs, such as prey by predators, affects the dynamics of ecological networks (e.g. plant-insects (1)), or the evolution of traits involved in interactions, as between hosts and their parasites (2). Decades of researches in ecology provided a vast catalog of possible factors affecting the variability of consumption rates within and across species, suggesting consumption rates are idiosyncratic and their variability is due to specificity at different scales, from the contingent evolutionary history of each species (3) to individuals’ personality, experience or ontogeny (4–7). Here, we challenge this view and test the hypothesis that observed within-species variations in consumption rates were mostly due to the intrinsic stochasticity of foraging in a depleted, unknown and spatialized environment.

Many factors have been identified as potential drivers of the variations of the consumption rates in predator-prey interactions, as testifies the abundant literature studying the evolution of traits and behaviors involved in foraging or the functional form of the relationship between prey abundance and predator consumption rates (the so-called functional responses). For instance, the time devoted to search for a prey depends on its local density (8–10), on the seasonal or spatial variation of the habitat (11, 12), the heterogeneity of prey distribution (13–15), the dimensionality of the environment (16–18), the presence of competing foragers (19) or the relative size between the prey and the predator (20, 21). Once a prey is found, the predator must spend some time to handle it which might depend on the quality of the prey (22) or on where and when it is more efficient to forage for another prey (23). Any other kind of interactions within or between species can also affect the consumption rates: predators can change their behavior in response to their own predators (24, 25), because of parasitism (26), if foraging is collective (27), or the rate at which prey are regenerated (28, 29).

A majority of studies have focused on a single factor among many others, in one or a few species. A few large scale analyses assessed whether one or several of these factors would drive variability of the consumption rates both within and between species. (30) showed that body mass and environment temperature generally affect the mean consumption rate across species. (31) focused on systematic statistical biases due to the non-linearity of the functions that are inferred in population ecology, casting doubt on estimations and models comparison, but also highlighting that experimental and methodological errors should be accounted for. All studies considered only three possible sources of variations (environmental, between-individuals or measurement errors), but did not consider variation that could come from the foraging process itself. Considering that foraging is a stochastic process in an unknown and depleted environment, the stochasticity of the foraging process itself is an unavoidable intrinsic source of variation of the consumption rate of prey by predators. Even if experimental conditions are perfectly controlled, if individuals are identical and if there are no measurements errors, foraging individuals would follow different paths when searching for prey, producing an intrinsic source of variation.

Our goal is to theoretically quantify this intrinsic source of variation and to compare it to experimental data. In the end, we aim at addressing the following questions: are observed variations of the same order of magnitude than the variations expected under stochastic foraging only? If so, that would suggest that environmental heterogeneity, between-individuals variability or measurements errors have negligible effect on a large scale on the variation of the consumption rates. Otherwise, if the observed variations are much larger than expected under foraging stochasticity only, that would be an evidence that these other sources of variability are the primary drivers. Finally, if observed variations are much smaller than expected, that would be evidence that some additional behavioral or ecological mechanisms are central. Our approach thus overall consists in providing predictions under a stochastic ’null’ or ’neutral’ model and evaluate to what extent it can explain the variability observed in data.

To this end, we derive the distribution of consumption rates that describes the number of prey consumed over a given foraging duration by a single predator modeled as a random walk, taking into account dimensionality, prey depletion, and handling and searching times. Although random walks are well-known objects extensively studied in mathematics and physics, their connection to consumption rates and functional responses in ecology has not yet been explored. We find sharp approximations of the distributions of the consumption rate by exploiting duality identities between a simple random walk and the foraging process. We then estimate the asymptotic of the range and the return times of a random walk, and we use uniform integrability via exponential moments to quantitatively predict the coefficient of variation, which is a standardized statistic measuring the relative magnitude of variations. We then compare the theoretical coefficient of variation of the consumption rates to the ones obtained in experimental and field observations. With numerical simulations, we then assess the generality or our results by estimating the robustness of the theoretical coefficient of variation. Our results suggest that the intrinsic stochasticity of the foraging process is the main driver of consumption rates variation, because predators visit already depleted sites a large proportion of their foraging time.

## Distribution of the consumption rate in a spatialized depleted environment

Our first goal was to quantitatively evaluate the order of magnitude of the variations of the consumption rates due to the stochasticity of the foraging process itself, while neglecting all other possible sources of variations (between-individual and environmental variabilities, measurements errors). We developed a stochastic model containing the simplest yet most fundamental mechanisms shared by predator species: predators forage in an unknown spatialized environment; they take some time to consume prey; their prey are depleted when consumed; and their total foraging duration is large relatively to moving from one site to another and handling a prey. We considered one predator following a symmetrical random walk on a regular grid, either in 1d, 2d or 3d, during a total duration *t*. Even though these assumptions are very simple, they are satisfied when the scale of movements are large enough relatively to body size and foraging bout duration (3). Each node initially contains a prey with probability *p*. The predator has no memory of its past foraging path and prey do not regenerate. The searching time *τ*_*e*_, i.e. the time taken by the predator to move from one site to another, only depends on the distance between sites. If the predator visits a site containing a prey, it spends there a time *τ*_*h*_ (the handling time) to consume the prey before moving to another site. If the site is empty, the predator randomly and immediately moves to one adjacent site. As the foraging path followed by the predator is stochastic, the total number of prey *Rt* consumed during the foraging bout with duration *t* is a random variable, as well as the consumption rate *Ft* := *Rt/t*. Assuming the duration of foraging *t* is large, the distribution of 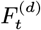 in a *d*-dimension space can be approximated by (proofs in Supp. Mat. A and B)

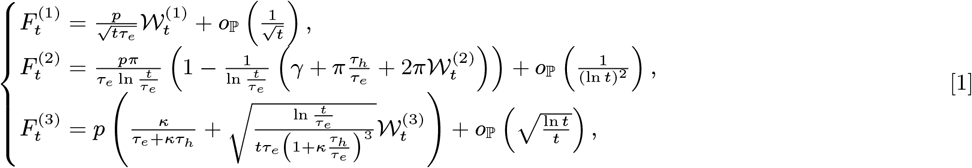

where *K, κ, γ* and *π* are numerical constant with known explicit values, 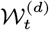 are random variables whose law only depends on dimension *d* = 1, 2, 3, *o*_ℙ_ (*η*_*t*_) means that the quantity is negligible compared to *η*_*t*_ in probability, *i*.*e*. for any *ε >* 0, ℙ (*o*_ℙ_ (*η*_*t*_) *≥ εη*_*t*_) *→* 0 as *t → ∞*. Eq. (1) shows that, depending both on the dimension of prey distribution in space and on the predator’s movements, the consumption rates follow different distributions 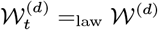. This is because the foraging path itself has very different properties: 𝒲^(1)^ is the difference between the maximal and minimal position of a 1-dimensional Brownian motion; 𝒲^(2)^is the local time of self-intersection of a 2-dimensional Brownian motion; 𝒲^(3)^ is a centered Gaussian distribution. Despite the simplicity of the process, only accounting for spatial structure, depletion, searching and handling, it gives rise to rich and non-trivial emerging properties: the form of the first order of the distribution approximation (the deterministic part) strongly depends on dimension: handling has no role in 1d, only a second order role in 2d, and a first order role in 3d. Eq. (1) also shows that the second order term, that reflects the magnitude of random fluctuations, can not be neglected in 1d or 2d as its magnitude is equal or similar to the first order term (the mean). In addition, stochastic fluctuations in 1d or 2d follow non-classical distributions. In a 3d environment, the random fluctuations of the consumption rate are expected to have a much smaller magnitude than the mean, and to follow a Gaussian distribution with a non-standard normalization.

## Coefficient of variation of the consumption rates due to foraging

Our goal is to evaluate the contribution of the intrinsic stochasticity of foraging on the within-species variation of consumption rates across all species and observational contexts. We thus chose the coefficient of variation as a standardized statistic for measuring the variability of the consumption rates. The expected coefficient of variation of the consumption rates under foraging stochasticity was calculated from the two first moments of the distribution of the consumption rate as 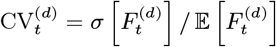, where 𝔼 and *σ* respectively are the mean and standard deviation. When the duration of foraging *t* is large, the coefficient of variation can be approximated for *t → ∞* by (Eq. (1), Supp. Mat. B)

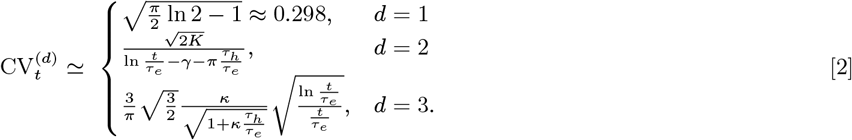

Our model predicts that the coefficient of variation of the consumption rate covers a very large range of orders of magnitude, but neither for all dimensions nor all parameter values (compare the green and red zones, and the gray line in Fig. 1). Roughly speaking (see also Mat. and Met.), the coefficient of variation should be of order 10^*−*1^ in 1d, at least of order 10^*−*1^ in 2d, and at most of order 10^*−*1^ in 3d. Under the assumptions that the searching time of sites with potential prey is much lower than the total foraging duration (*t/τ*_*e*_*≫* 1, e.g. (24, 32)) and that the handing time is at most of the same order than the searching time (*τ*_*h*_*/τ*_*e*_ *∼ O*(1), e.g. (24, 30, 32)), the coefficient of variation when considering the stochasticity of foraging only, can even further be expected to be almost constant and lie between orders 10^*−*1^ and 1 in 1d and 2d, or to be very small (*≪*10^*−*1^) in 3d (Fig. 1).

**Fig. 1.**
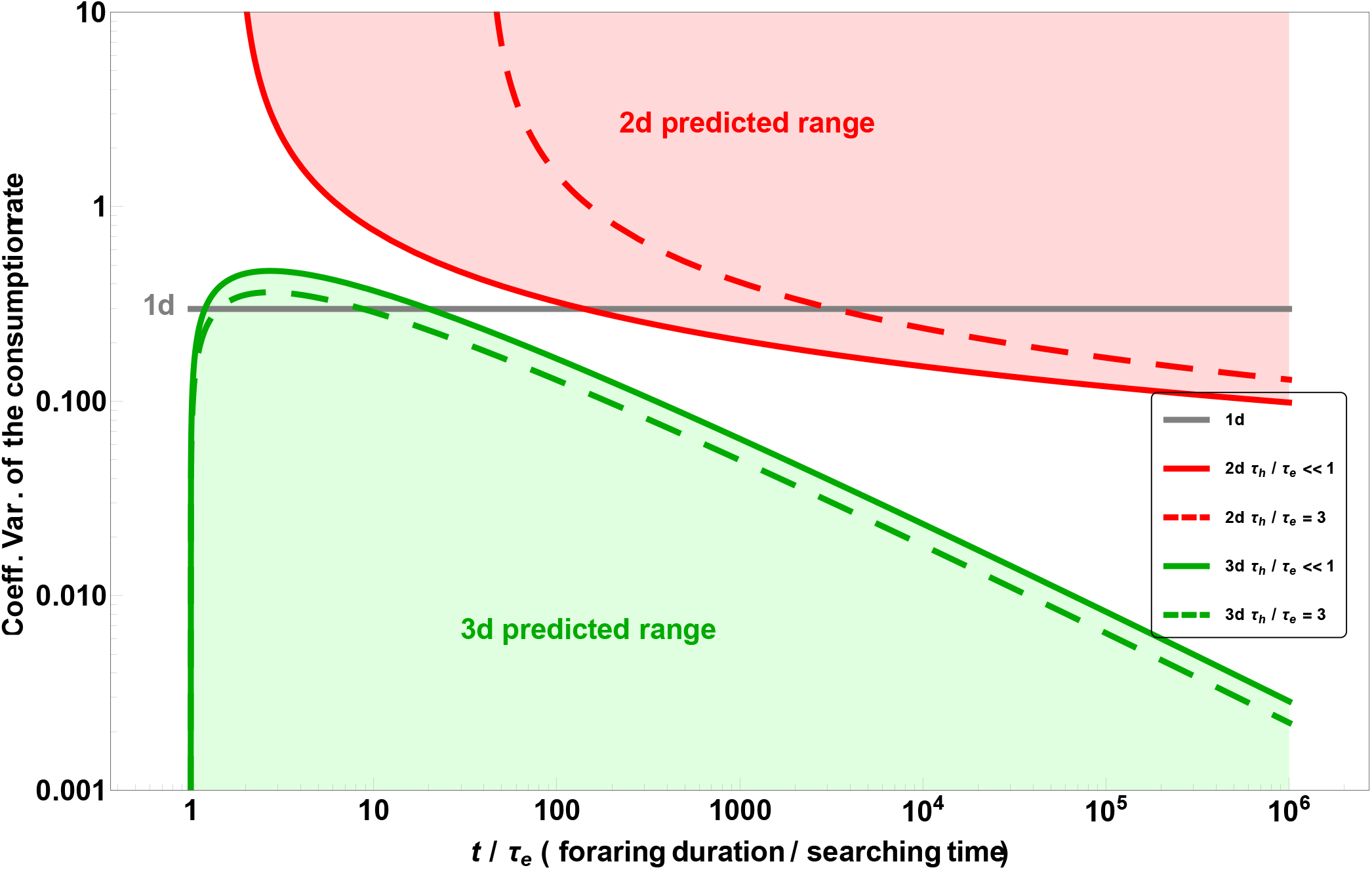
Predicted range of the coefficient of variation of the consumption rate in a simple random walk with handling (Eq. (2)). Solid lines (no handling time, *τ*_*h*_ = 0): minimum coefficient of variation in 2d (red solid line); maximum coefficient of variation in 3d (green solid line). Solid gray line: coefficient of variation in 1d (independent of the total foraging time *t*, foraging time between sites *τ*_*e*_ and handling time *τ*_*h*_).

## Predicted vs. observed coefficients of variation

In order to test to what extent the stochasticity of foraging on its own drives the variability of the consumption rates, we compared our model’s quantitative predictions to coefficients of variation estimated from data. The consumption rates were collected from three categories of datasets, with different uncertainty levels and experimental controls, and different potential sources of variations (see Methods and Supp. Mat. E for further details). Dataset 1: the highest level of uncertainty as it largely comes from the automatic digitization from figures and tables in published papers. Dataset 2: an intermediate level of uncertainty as it was not known whether a single individual was used different times for a given or different treatments, which made not possible the distinction between within and between-individuals variability. Dataset 3: the lowest uncertainty as the observed individuals were known and observed several times. We then calculated the coefficient of variation of the foraging rates for all datasets for each experimental treatments (i.e. generally for an environmental condition, a given pair of prey and predator species, and a given initial prey density). We obtained more than 3800 estimations of the coefficient of variation, for more than one hundred species of prey and one hundred species of predators, for species ranging from unicellular (e.g. ciliates) to vertebrate (e.g. fish and birds), for initial prey density varying by up to 15 orders of magnitude across experiments. We then compared the range of observed values with the expected values under foraging stochasticity only (Fig. 2, Fig. Supp E.1). For all three datasets, a large majority of the measured coefficients of variation lie between 0.1 and 1 (respectively 73.2%, 87.8% and 85.1% for datasets 1, 2 and 3; Fig. 2(a-c), Fig. Supp E.1). The medians are close to 0.298, the value predicted in 1d (respectively 0.291, 0.408 and 0.273). In a large majority of cases, the observed coefficients of variation falls within the range predicted in our model in a 1d or 2d environment. In addition, the predicted range of the coefficients of variation in 2d is in line with a larger proportion of observations than in 3d for all three datasets (85.2% vs. 69.3%, 92.3% vs. 60.3%, and 86.9% vs. 75.5%, 2d vs. 3d, in datasets 1, 2 and 3, respectively). It suggests that foraging effectively occurred as in a 1d or 2d space. Figure 2(a-b) also shows that the dimensionality of the experiment has no effect on the range of the observed coefficients of variation, as the observed coefficients of variation cover the whole range of predicted values (compare colored dots in Fig. 2(a) and (b)).

**Fig. 2.**
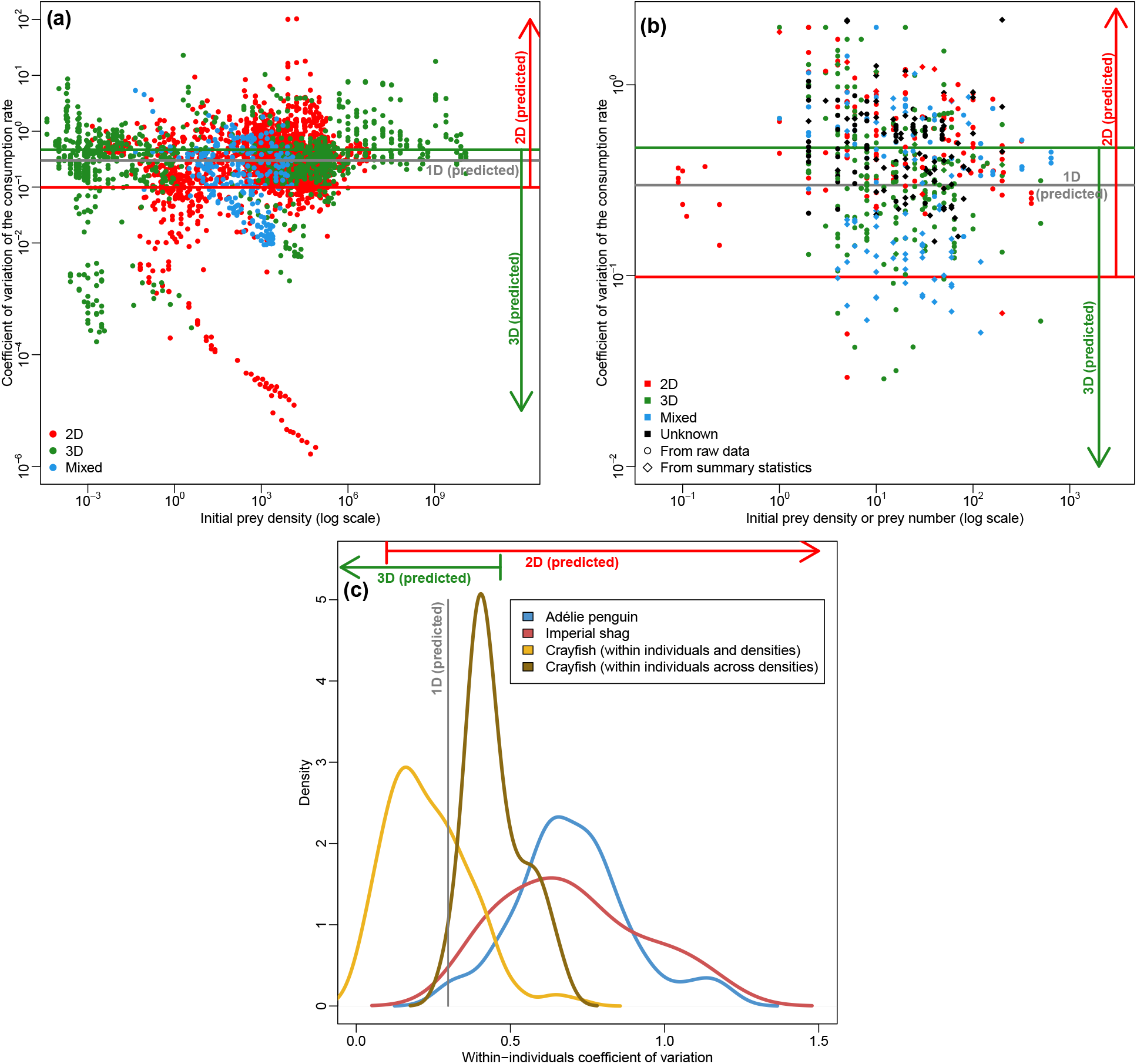
Coefficients of variation of prey consumption rates in experiments compared to the maximal range predicted by the model (lines). (a) Dataset 1: data from the FoRAGE database; (b) Raw data (between and within individuals variation confounded): (c) Raw data (between and within individuals variation isolated). Dimensionality of space in the model and experiment is represented by colors: Models in 1d, 2d, 3d respectively in black, red and green; Experiments in 2d, 3d or mixed (prey distributed on surfaces in a 3d system, e.g. on leaves in a terrarium) respectively in red, green and blue.

We then assessed the validity and robustness of the predicted coefficient of variation using numerical stochastic simulations by accounting for additional ecological mechanisms possibly adopted by different species: memory, preferred foraging direction and larger movements. Fig. 3 (a) shows that the coefficient of variation predicted in our model is very close to simulations in 1d, but it is slightly overestimated in 2d and 3d (predicted and simulated values are yet of similar order of magnitude). This overestimation might be due to a very slow speed of convergence of the approximation as the predicted values slightly gets closer to simulations when times *t* increases (Fig. 3 (a)). As predicted by our model (Eq. (2)), the probability *p* that a site initially contains a prey does not affect the coefficient of variation (Fig. 3(b)). Short term memory slightly decreases the coefficient of variation (compare Figs. 3(a) and (c)). Longer movements significantly decrease the coefficient of variation in 2d but only when the jump range is very large (dark dots), otherwise they do not affect it in 1d and only slightly in 3d (Fig. 3(d)). Finally, the coefficient of variation is strongly affected when the predator has a preferred foraging direction, which is not surprising as the foraging path becomes more deterministic as the weight of direction preference increases (Fig. 3(e)). Overall, we found that our model is robust to additional mechanisms except in situations where the stochasticity of the foraging paths is decreased. This is consistent with general properties of random walks (see Supp. Mat. A4). Hence, additional mechanisms can, on the one hand, have little effect on the coefficient of variation which would explain why most observed values fall in the range predicted by our model (between 0.1 and 1). On the other hand, they can partly explain why observed coefficients of variation are much lower than 0.1.

**Fig. 3.**
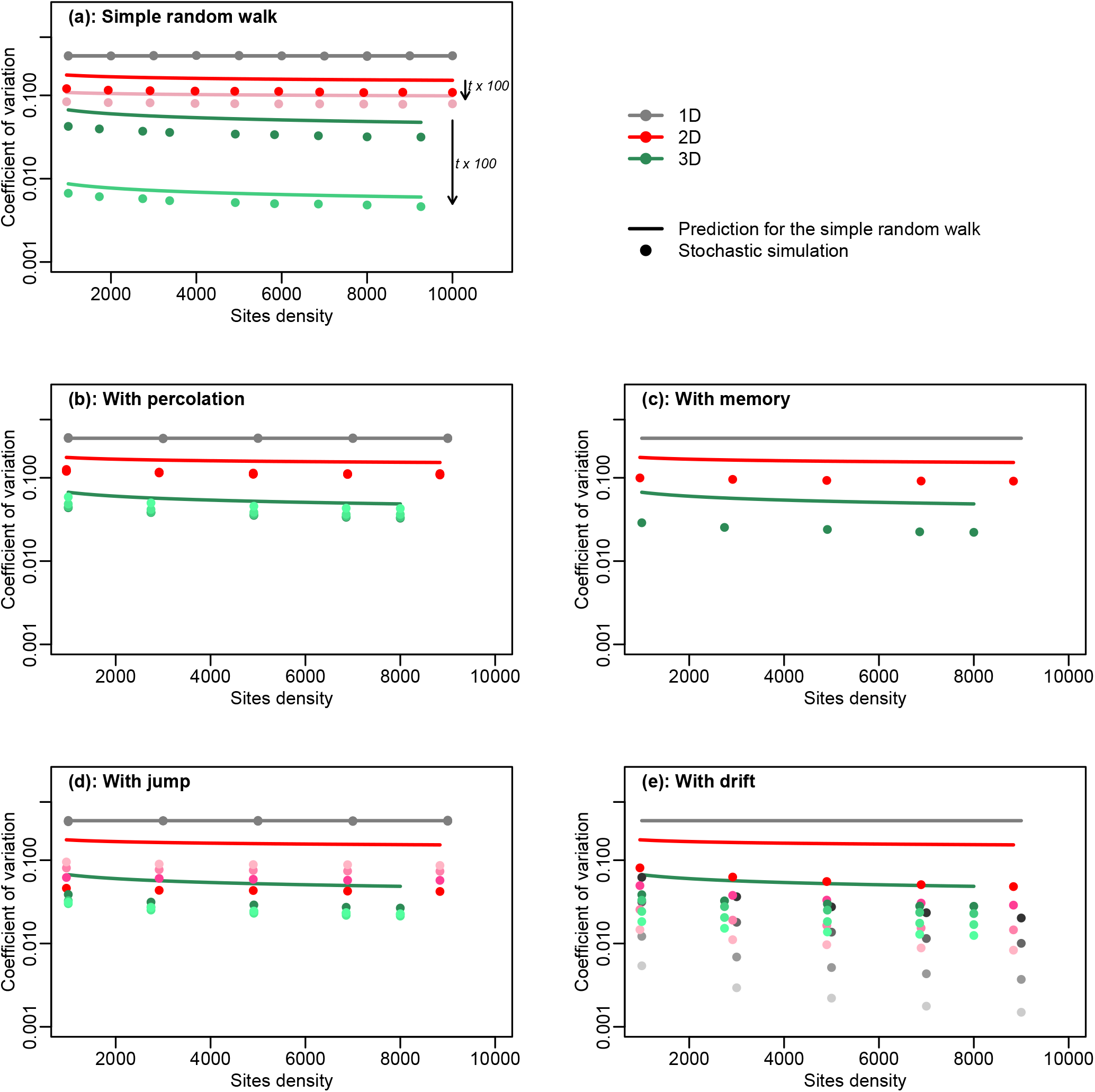
Robustness of the prediction of the coefficients of variation. The sub-figures compare the value predicted of the coefficient of variation in 1d, 2d or 3d from Eq. (2) (solid line) and the value estimated from exact stochastic simulations (dots) where additional mechanisms are added to the reference model (a) (Eq. (2)) with sites initially containing a prey with probability *p* = 0.5 for two total foraging times *t* = 10^5^ and *t* = 10^7^; Sub-figure (b) shows coefficients of variation where sites initially contained prey with probability *p* = 0.05, 0.1, 0.25, 0.5 (from darker to lighter colors); (c) With short term memory (the predator avoids the immediate previously visited site); (d) With jump: The predator can move to more distant site in one step. The distance follows a power-law distribution with exponent *θ* = 2, 2.5, 3, 3.5 (from lighter to darker colors, shorter to larger jumps); (e) With drift: The predator has a preferred foraging direction with weight *µ* = 0.05, 0.1, 0.25, 0.5 (from darker to lighter colors). Default values for simulations (unless indicated): *t* = 10^7^, *τ*_*h*_ = 0.1, *τ*_*e*_ = *L/*(*x*^1*/d*^ *−* 1) with *x* the sites density and *L* = 10^3^ an arbitrary length value between two sites.

## Searching for prey is the main driver of consumption rate variability

In summary, despite several sources of variations in experiments could contribute to the estimated coefficient of variations, stochasticity of the foraging process alone is quantitatively compatible with the range covered by data. Two possibilities can explain that the coefficients of variation estimated from observations mostly lie between 0.1 and 1 across a large majority of species and observational contexts (Fig. 2, Supp. Inf. C)). *Hypothesis 1* : Foraging paths stochasticity is negligible relatively to other sources of variability. If so, one should explain how is it possible that between-individual variability, environment heterogeneity and measurements altogether scale such that the estimated coefficients of variation mostly varies in such a small range of values between (0.1-1) across all species and experimental contexts. *Hypothesis 2* : Foraging paths stochasticity is the main driver, and the effective dimension of foraging is 1d or 2d. As our model’s predictions are in line with most of the estimated coefficients of variations, it suggests that experimental errors, environmental and between-individuals variabilities are negligible. In other words, the coefficient of variation of the consumption rate is large in 1d and 2d because the randomness of foraging paths is strongly linked to the randomness of the number of prey effectively consumed. In 3d, paths are also highly random but the number of prey consumed is very little affected by the particular trajectory taken by the predator.

Between-individual, between-species, or environmental variabilities can still affect the consumption rates as its mean and variance depend on the handling and searching times (Eqs. (3) and (4)):

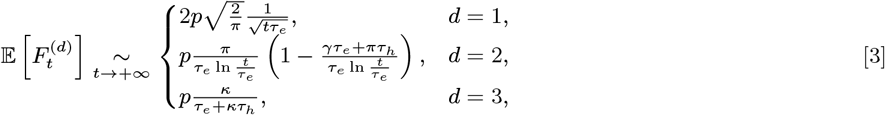

and

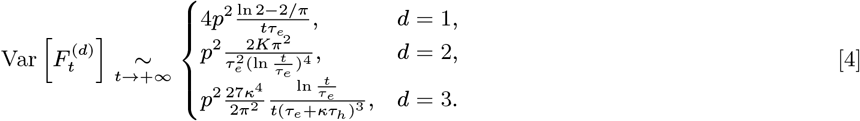

However, the average of the consumption rate and the magnitude of its fluctuations, measured by the standard deviation, both similarly depend on the parameters of the model in 1d and 2d, in such a way that they are of the same order. In 3d, standard deviations are on the contrary expected to be much smaller than the average. That is why the coefficient of variation is expected to be large for any species or ecological contexts.

## Dimensionality matters because of depletion

The mean and variance of 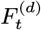 are very different depending on dimension (Eq. (3) and Eq. (4)). The relative importance of the searching vs. handling times in a functional response with depletion depends on dimensionality not because of the predator-prey interaction itself (as suggested in (16)), but because the time spent to successfully forage non-empty sites grows on different scales. In 1d, the handling time *τ*_*h*_ is negligible, while in 3d the searching and handling times have an effect of the same order. In 2d, handling has an effect of the same order than searching when foraging duration is short, otherwise it has a second order effect on the consumption rate. Consequently, estimating a parameter such as the handling time would need controlling for the time scale of depletion relatively to foraging duration in experiments. Eq. (3) also shows that the decrease of the consumption rate with foraging duration *t* depends on dimension: it decreases rapidly in 1d (as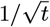), slowly in 2d (as 1*/* ln *t*) and is constant in 3d (as 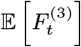 does not depend on *t* at first order).

Eq. (4) shows that handling time has no effect on the variance of the consumption rate 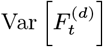 in a 1d or 2d environment, even if the repartition of prey in the environment is initially homogeneous (*p* = 1). In contrast, both handling and searching times equivalently affect it in 3d. It thus suggests that handling would little contribute to variation of the consumption rates in a 1d or 2d depleted environment. Eq. (4) also shows that the size of the fluctuations are differently affected by the foraging duration *t*: in 1d, fluctuations are expected to have the same order as the mean and standard deviation both decreases as 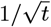 in 2d, fluctuations are expected to be lower than the mean but of similar order, (the mean decreases as 1*/* ln *t* and the standard deviation as 1*/*(ln *t*)^2^; in 3d, fluctuations are expected to be of an order lower than the mean (as the mean does not depend on *t* while the standard deviation rapidly decreases as 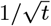). This means that it is expected that variability of consumption rates should be negligible in a 3d environment when compared to the mean, but at least of similar order of magnitude in a 1d or 2d environment.

## Bridging the gap between foraging theory and functional responses in ecology

Assuming that the searching time *τ*_*e*_ is a decreasing function of prey density *x* (*e*.*g*. 1*/x*^(1*/d*)^), our model also provides functional responses, i.e. a function describing how consumption rate of a predator is related to prey density (8)(Fig. Supp. C.1). Many functional responses have been proposed under the assumption that prey depletion is negligible (33–35), only a few with prey depletion (29, 36, 37). All functional response models have neglected space, the foraging process and its stochastic nature, thus perpetuating a division between the fields interested either in foraging or functional responses (see (9) for a critical review). Here, by explicitly including the foraging process of prey distributed in space, even though under simplifying assumptions, we contribute in bridging the gap between those two mostly independent fields with promising outcomes. For instance, the form of the functional responses with depletion including the foraging process are very different than the one which is mostly used, the Rogers-Royama equation (29, 36, 37). We also show that in a 3d environment, the first order approximation gives the form of a Holling type II functional response. This suggests that depletion would have little effect on the functional response in cases where prey are distributed in 3d. Finally, as our results show that fluctuations of the consumption rate are of the same order than the average, it suggests that using deterministic models such as the Rogers-Royama equation for interpreting data or estimating parameters is limited.

Another assumption differs between the Rogers-Royama model with prey depletion and ours: they supposed a fixed initial number of prey while we supposed an open-ended environment with virtually no prey limitation. At first sight, supposing a fixed initial number of prey would be closer to experimental design as prey are generally distributed in a closed environment at the start of the experiments. However, data show little relationship between initial density of prey and the coefficient of variation (Fig. 2), as predicted by our model (Eq. (2), Fig. 3). In fact, we collected the coefficients of variation from data only when there was variation in the number of prey consumed, i.e. when *CT >* 0. Cases where CV = 0 were found in data only when the initial number of prey was low, i.e. when for a given initial density all prey were consumed by each predator in all replicates. As the order of magnitude of the estimated coefficient of variation lie within the range predicted by our model, it suggests that as soon as there exists some variation in the number of prey consumed, the prey-predator system works as if in an open-ended space. This should depict experimental conditions where the total duration of the experiment was short enough that prey were not fully depleted, but long enough relatively to the handling and searching times. Finally, despite we do not impose a fixed initial number of prey, the order of magnitude of the variation of the consumption rates is well captured by our model, further strengthening our conclusion that the stochasticity of the foraging paths is the main driver.

## Theoretical and empirical implications

The fact that foraging variations are mostly driven by foraging paths stochasticity suggests that the handling time has a negligible effect on consumption rate relative to searching time. This is true for both the coefficient of variation and the expectation (Eq. (3)). It also suggests that the variability of the consumption rate within and between species is little affected by the individuals or species properties such as variation in size or other traits, or trophic levels, and that foraging occurs in an environment with an effective dimension 1d or 2d, even when it is actually in 3d. This can be an evidence that predators actually forage in 1d or 2d (38, 39), or, not exclusively, that prey are distributed in 1d or 2d. Finally, other mechanisms certainly affect the consumption rates as many papers showed how animals tend to change their behavior depending on the prey, habitat, their own state, etc. However, our results show that additional mechanisms have negligible impact on within-species variability compared to the intrinsic variability due to searching for prey.

As the predictions from our model are mostly compatible with data, stochastic foraging in a spatialized and depleted environment could be seen as a null model for consumption rates and functional responses studies. It is important because many studies aim at inferring parameters such as the attack rate from experimental data (24, 29, 30). As the variation of the consumption rate has the same order of magnitude than the mean, it makes inference difficult: the control one might have on the quality of an estimator is limited by the inherent noise from the observed process and its outcomes. This result is also important from both a theoretical and empirical viewpoints, as stochasticity in population biology is generally thought to be important only when population is low (e.g. genetic drift, demographic stochasticity). Our results show that it is generally not the case as stochasticity is expected to be large even in large systems on a short time scale. The resulting implications of the interactions stochasticity on the understanding and modeling of populations and communities dynamics remains an open question. On a side note, our results provide a ’quick and dirty’ criterion for evaluating the validity of a stochastic model of foraging or prey consumption of a predator: its coefficient of variation should lie between 10^*−*1^ and 1, and close to 0.3 on average.

As a null model, our approach can also help explaining the observed discrepancies between data and predictions (Fig. 2) such as why coefficients of variation range from one to (rarely) several orders of magnitude lower than 10^*−*1^. Our numerical simulations suggest that some non-exclusive additional mechanisms, such that predator’s memory or a preferred foraging direction, can significantly decrease the coefficient of variation. Our model also suggests that in some experimental contexts, the effective dimension of foraging is closer to 3d, or that it could be due to the fast regeneration of prey(34), which would however be inconsistent with most experimental conditions. Finally, it could be due to experimental design itself as the boundaries of the environment can be either attractive (39) or repulsive. Coefficients of variation which range from one to several orders of magnitude larger than 1 can be an evidence that other sources of variation are at least as important as the intrinsic stochasticity of the foraging paths. It can also be due to particular situations in 2d such as a large searching or handling times compared to the total foraging duration (Fig. 1).

Finally, our results have implications for the evolution of foraging and the study of the optimal foraging theory. The fitness of individuals depends on the variance of the parameters of the system (40) but also on the inherent variance of the process itself as shown here. As this variance depends on dimensionality, foraging strategies can be expected to have differently evolved depending on dimensionality, yet independently of spatial heterogeneity. Using simulations of random walk in 2d, (41) predicted that optimal foraging strategies would evolve directional motion (i.e. movements with little turning). However, we showed that a preferred direction for foraging tends to largely decrease the coefficients of variation, which would not be consistent with data. This would surprisingly suggest that species have evolved sub-optimal foraging behaviors, as the observed coefficients of variations are larger than expected under a preferred foraging direction. Otherwise, it more likely suggests that other traits and mechanisms have evolved in order to optimize foraging efficiency in an unknown spatialized environment rather than a preferred direction.

### Supporting Information (SI)

Supporting information are provided as a separate file.

***SI Datasets***. Information about data sources and accessibility are given in Supp. Inf. E.

## Materials and Methods

Please describe your materials and methods here. This can be more than one paragraph, and may contain subsections and equations as required.

### Model

A forager performs a random walk in a *d*-dimension environment where prey are distributed on a lattice. The time taken to go from one site to another is *τ*_*e*_. Note that by assuming that the searching time *τ*_*e*_ is proportional to *x*^(*−*1*/d*)^, where *x* is the density of sites potentially containing prey, our functions *F* ^(*d*)^ are functional responses, as classically defined. If a site is visited for the first time, it contains a prey with probability *p*. Consuming the prey takes a handling time *τ*_*h*_. When a prey is consumed, it is not renewed (the time scale for prey renewal is supposed much larger than the foraging time scale). Given these times, the distribution of *Rt*, the number of prey consumed after a time *t*, is linked by a duality argument to the distribution of the minimum number of searching steps needed to consume at least a given number of prey. By exploiting the convergence in law of the distribution of the number of distinct sites visited after a given number of steps (42–44), the distribution of 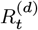 converges in law as *t → ∞* to different explicit expressions depending on the dimensionality of the environment (Supp. Mat B). Approximations of the first two moments of 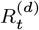 are then obtained. Finally, the distribution of the consumption rates and its first two moments are then deduced as 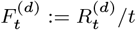 (see details in Supp Mat. A and B).

### Orders of magnitude of the coefficient of variation

Depending on the dimension, the coefficient of variation is expected to differ in orders of magnitude (Fig. 1, note the log-scale on the *y*-axis). When the total foraging time *t* is large enough, the coefficients of variation in a 1d, 2d, 3d depleted environments are respectively expected to be of order 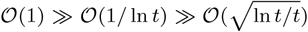. Note that the order of magnitude of the coefficient of variation in a renewed environment is close to the one in a 3d depleted environment: 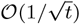 (34). In other words, in 1d, the coefficient of variation is expected to be constant. As ln *t/τ*_*e*_ increases very slowly with *t/τ*_*e*_, the coefficient of variation in 2d is expected to be of an order of magnitude either slightly lower or larger than in 1d (compare the red zone and the gray line in Fig. 1). On the contrary, the coefficient of variation in 3d is expected to rapidly decrease with *t/τ*_*e*_ and thus to be at least one order of magnitude lower than in 1d and 2d. This is because stochastic fluctuations of the number of prey consumed are generally expected to be much larger in 1d or 2d than in 3d. Eq. (2) also shows how the coefficient of variation of the consumption rate is expected to be affected by the ecological and behavioral parameters. In 1d, the coefficient of variation does not depend on any parameter. In 2d and 3d, the coefficient of variation is not affected by the initial heterogeneity of prey repartition *p*, while it is affected by the searching and handling times *τ*_*e*_ and *τ*_*h*_ in different ways. An increasing searching time *τ*_*e*_ generally increases the coefficient of variation in 2d and 3d, but has a non-monotonous effect in 3d as it decreases it for low values of *t/τ*_*e*_. The handling time has opposite effect on the coefficient of variation as it increases it in 2d while it decreases it in 3d (Eq. (2), compare full and dashed lines in Fig. 1). In particular, the coefficient of variation shows a maximal value for *τ*_*h*_ = 0 in 3d, while it can take arbitrarily high values in 2d.

### Stochastic numerical simulations of foraging rates

The path followed by a foraging individual was numerically simulated on a *d*-dimensions lattice where each node initially contains a prey with probability *p*. The individual moves to the next site, which is randomly chosen with equiprobability among the four nearest (cardinal) ones. The sites are homogeneously distributed with a density *x* on the lattice. One thousand stochastic simulations are generated for all set of parameters values (sites densities *x* and total foraging duration *t*). The empirical distribution of the foraging rates and its expectation were estimated. In order to test for the robustness of the approximations obtained with our model, we added behavioral features: random walk with (stochastic) jumps to distant sites, with preferential direction (drifted random walk), with short-term memory (avoidance of the last visited site).

### Datasets

The coefficient of variation of the consumption rate were estimated from three different datasets with different degree of uncertainties regarding the sources of variability (*i*.*e*. data collection methodology, experimental controls, between-individuals variability, within-individuals variability, controlled or uncontrolled environmental variability, variability due to the foraging process itself). Further details are given in Supp. Mat. E. <u>Dataset 1:</u> We extracted relevant data from the FoRAGE database (45) which compiles functional responses from controlled experiments in the published literature. FoRAGE database was populated by raw data or summary statistics either automatically digitized from articles’ figures or gathered from articles’ tables. There were often uncertainties about how many actual observations were represented by one dot on figures (several dots overlapped if individuals in a replicate consumed the same number of prey). Another uncertainty source was the standard deviations with zero values, which can either be due to entries errors in the database or because of a true absence of variation. We thus decided to discard all experiments with no variation within an experimental treatment in the database. In order to be conservative and to limit noise due to data collection, we also discarded experiments from the FoRAGE database that do not satisfy the following criteria: (i) at least eight independent measurements for a given experimental treatment and (ii) at least 80% of raw data recognizable from the figures among all replicates for a given treatment. We overall estimated 3181 coefficients of variation for various experimental conditions (regarding prey density, temperature, environmental dimensions, prey and predator species, etc.), for 126 predator and 104 prey species, covering a biomass range of near 7 and 12 orders of magnitude, respectively. We sorted data regarding the dimension of the environment (as acknowledged by the database’ authors): 2d, 3d, or a mix between 2d and 3d (for instance when prey were distributed on plant leaves within a 3d space such as a terrarium). Overall, the coefficient of variation calculated from Dataset 1 may come from different sources of variation: measurement errors including populating the database and digitization, between and within individuals variability, the stochastic foraging process itself, controlled or uncontrolled environmental variability. <u>Dataset 2:</u> We estimated 602 coefficients of variation from 41 different raw datasets collected either from the Dryad repository, or the database by (31), or directly from the authors. We only considered datasets with a single species of prey in a given experiment, with known initial prey densities or number, and with replicates within densities. The coefficients of variations were directly calculated from raw data when available, or from the mean, the standard error and the number of replicates otherwise. It was generally not known whether replicates used the same individuals several times independently or not. When the experiments considered different treatments for a given density (e.g. different temperatures or prey size), we calculated the coefficient of variation across treatments. Overall, the coefficient of variation calculated from Dataset 2 potentially comes from different sources of variation: measurement errors, uncontrolled environment variability, between and within individuals variability, or the stochastic foraging process itself. <u>Datasets 3:</u> We estimated 229 within-individuals coefficients of variation from datasets kindly provided by Yuuki Watanabe, Agustina Gómez-Laich and Stefan Linzmaier. In the three cases, several measurements were obtained from the same individuals thus allowing to isolate the within individuals from between-individuals variability. These datasets included the consumption rates observed for several individuals during different sessions of observations for each individual. Overall, the coefficients of variation estimated from Dataset 3 potentially comes from different sources of variation: measurements errors, within individual variability, uncontrolled environmental conditions, and the stochastic foraging process itself.

## ACKNOWLEDGMENTS

Kilian Raschel for his help in literature searching for fine estimates of return times to the origin of random walks, Eliza Vergu for helpful discussions, Yuuki Watanabe and Stefan Linzmaier for sharing their raw datasets, Agustina Gómez-Laich for having provided the coefficient of variation from their dataset. The authors acknowledge the support of the “Chaire Modélisation Mathématique et Biodiversité (MMB)” of Veolia Environnement-Ecole Polytechnique-Museum National d’Histoire Naturelle-Fondation X, and of the grant MITI CNRS 80’.

