## Supplementary information for "Stochastic foraging paths primarily drives within-species variations of prey consumption rates"

Vincent Bansaye[a], Geoffroy Berthelot[a,b,c], Amina El Bachari[a], Jean-René Chazottes[d], and Sylvain Billiard[e,1]

[a]Centre de Mathématiques Appliquées (CMAP), CNRS, Ecole polytechnique, Institut Polytechnique de Paris, Palaiseau, France

[b]Institut National du Sport, de l'Expertise et de la Performance (INSEP), Paris 75012, France

[c]Research Laboratory for Interdisciplinary Studies (RELAIS), Paris 75012, France

[d]Centre de Physique Théorique, CNRS, Ecole polytechnique, Institut Polytechnique de Paris, Palaiseau, France

[e]Univ. Lille, CNRS, UMR 8198 – Evo-Eco-Paleo, F-59000 Lille, France

May 29, 2024

### A Behaviors of the range of random walks

We first gather various results on the number of distinct sites visited by a simple, symmetric, random walk on $\mathbb{Z}^d$. This random variable is usually called the "range" of the random walk. It is the central quantity to define the "functional response" in our context. We will state known results for any dimension $d$, with a particular focus on dimensions $d = 1, 2, 3$. It turns out that in $d = 1$ and $d = 2$ the mean number of distinct visited sites grows more slowly than the duration $n$ of the random walk, a consequence of "oversampling" as each site is repeatedly visited while the number of visits grows with $n$. We in particular show that, even in the simplest situation, rich and diverse behaviors of the random walks emerge according to $d$. We will indicate below the degree of generality in which the following results remain valid (with slight modifications).

#### A.1 Setting and two basic results

Let $d \geq 1$ be an integer and denote by $(\mathrm{e}^{(1)}, \ldots, \mathrm{e}^{(d)})$ the canonical basis for the lattice $\mathbb{Z}^d$, so we have $i = (i^{(1)}, \ldots, i^{(d)})$ for $i \in \mathbb{Z}^d$. We call "sites" the elements of $\mathbb{Z}^d$. We focus on $d = 1, 2, 3$. Let $(X_n)_{n \geq 0}$ be the simple symmetric random walk on $\mathbb{Z}^d$ starting from the origin, where $X_n = \big(X_n^{(1)}, \ldots, X_n^{(d)}\big)$ is the position of the random walker at time $n$ on $\mathbb{Z}^d$. This process can be viewed as a Markov chain with state space $\mathbb{Z}^d$, initial state $X_0 = 0 := (0, \ldots, 0)$, and transitions probabilities (for $n \geq 0$)

$$\mathbb{P}\big(X_{n+1} = j \,|X_n = i\big) = \frac{1}{2d}$$

if $j - i \in \{\pm\,\mathrm{e}^{(1)}, \ldots, \pm\,\mathrm{e}^{(d)}\}$ (that is, if $i$ and $j$ are nearest neighbors), and this transition probability is equal to 0 otherwise. The process $(X_n)_{n \geq 0}$ can also be seen as a sum of a sequence of independent identically distributed random variables. Letting $(\xi_k)_{k \geq 1}$ be a sequence of i.i.d. random variables taking values in $\{\pm\,\mathrm{e}^{(1)}, \ldots, \pm\,\mathrm{e}^{(d)}\}$ such that

$$\mathbb{P}\big(\xi_k = \pm\,\mathrm{e}^{(i)}\big) = \frac{1}{2d}, \quad i = 1, \ldots, d$$

we define

$$X_0 = 0 \quad \text{and} \quad X_i = \sum_{k=1}^{i} \xi_k, \quad i = 1, 2, \ldots.$$

**Definition A.1.** *The range of the random walk is the sequence of random variables defined by*

$$\mathtt{N}_n = \mathrm{Card}\{X_i : i \leq n\} = \mathrm{Card}\big\{k \in \mathbb{Z}^d : \exists\, i \leq n \textit{ such that } X_i = k\big\},\ n \geq 1.$$

*By convention we set* $\mathtt{N}_0 = 1$.

Hence, $\mathtt{N}_n$ is the number of distinct sites visited up to time $n$ by the random walk. Note in passing that $(\mathtt{N}_n)_{n \geq 0}$ is a non-Markovian process.

Let

$$\kappa = \kappa(d) = \mathbb{P}(\text{no return to } 0) = \mathbb{P}(\forall n > 0 : X_n \neq 0). \tag{A.1}$$

The first result about the range is a kind of "law of large numbers".

**Theorem A.2.** *Let* $d \geq 1$. *Then*

$$\mathbb{P}\left(\frac{\mathtt{N}_n}{n} \xrightarrow[n\to+\infty]{} \kappa(d)\right) = 1. \tag{A.2}$$

It is well known that when $d = 1$ or 2, the walk is recurrent, that is, it visits any site infinitely many times with probability one, hence, in particular, $\kappa(1) = \kappa(2) = 0$. When $d \geq 3$, the random walk is transient, hence $0 < \kappa(3) < 1$. See [1, p. 38] for a proof.

The second basic result about the range tells us that, when $d \geq 2$, $\mathtt{N}_n \underset{n\to+\infty}{\sim} \mathbb{E}(\mathtt{N}_n)$, with probability one.

**Theorem A.3.** *Let* $d \geq 2$. *Then*

$$\mathbb{P}\left(\frac{\mathtt{N}_n}{\mathbb{E}(\mathtt{N}_n)} \xrightarrow[n\to+\infty]{} 1\right) = 1. \tag{A.3}$$

See [2] for a proof. Notice that this result is not true for $d = 1$. In that case, we will see that $\mathtt{N}_n / \mathbb{E}(\mathtt{N}_n)$ converges only in law (to some non-degenerate random variable).

#### A.2 Link with return times to the origin

Let $d \geq 1$. For $j \geq 1$, we define new random variables $\Upsilon_j$ associated to the random walk by

$$\Upsilon_j = \begin{cases} 1 & \text{if the } j\text{-th step hits a new site} \\ 0 & \text{otherwise.} \end{cases}$$

Formally, $\Upsilon_j = \mathbb{1}_{\{X_j \neq X_{j-1}, X_j \neq X_{j-2}, \dots, X_j \neq X_0 = 0\}}$. Hence we have

$$\mathtt{N}_n = \sum_{j=1}^{n} \Upsilon_j \tag{A.4}$$

since we are exactly counting the total number of distinct visited sites when the random walk made $n$ steps. Next, for $j \geq 2$,

$$\mathbb{E}(\Upsilon_j) = \mathbb{P}\left(X_j \neq X_i \text{ for } i = 1, \dots, j-1\right)$$

(we have $\mathbb{E}(\Upsilon_1) = \mathbb{P}(X_1 \neq X_0) = 1$), that is, $\mathbb{E}(\Upsilon_j)$ is the probability that the $j$-th step takes the walk to a non-visited site in any of the preceding $j - 1$ steps. The basic but fundamental observation is that

$$\mathbb{E}(\Upsilon_j) = \mathbb{P}(T_0 > j) = \sum_{k=j+1}^{+\infty} \mathbb{P}(T_0 = k) \tag{A.5}$$

where $T_0 = \inf\{n \geq 1 : X_n = 0\}$, the first time the walk hits the origin. That is, the expected value of $\Upsilon_j$ is exactly the probability that the walk did not come back to the origin in $j$ steps (since it starts at the origin). Hence, in view of (A.4) and (A.5), we have

$$\mathbb{E}(\mathtt{N}_n) = \sum_{j=1}^{n} \mathbb{E}(\Upsilon_j) = \sum_{j=1}^{n} \mathbb{P}(T_0 > j) = \sum_{j=1}^{n} \sum_{k=j+1}^{+\infty} \mathbb{P}(T_0 = k). \tag{A.6}$$

Hence, we are left to estimate $\mathbb{P}(T_0 = k)$ $(k \in 2\mathbb{N})$ to estimate $\mathbb{E}(\mathtt{N}_n)$. [1]

[1] We let $\mathbb{Z}_+ = \{0, 1, 2, \dots\}$ be the positive integers including 0, and we will denote by $\mathbb{N} = \{1, 2, \dots\}$ the set of strictly positive integers.

**Remark A.4**. *Observe that* $\kappa = \lim_{j\to+\infty} \mathbb{P}(T_0 > j) = \lim_{j\to+\infty} \mathbb{E}(\Upsilon_j)$ *(see* (A.1)*).*

**Remark A.5**. *It turns out that* (A.5) *(hence* (A.6)*) is valid for any random walk because it only relies on the fact that the random variables* $\xi_k$ *are independent and identically distributed, which implies that* $(\xi_1, \xi_2, \ldots, \xi_j) \stackrel{\text{law}}{=} (\xi_j, \xi_{j-1}, \ldots, \xi_1)$*, for all* $j \geq 1$*. Hence, we can "reverse" (in law) the path of the random walk to get*

$$\begin{aligned}\mathbb{P}(X_j \neq X_{j-1}, X_j \neq X_{j-2}, \ldots, X_j \neq X_0 = 0) &= \mathbb{P}(\xi_j \neq 0, \xi_j + \xi_{j-1} \neq 0, \ldots, \xi_j + \xi_{j-1} + \cdots + \xi_1 \neq 0)\\ &= \mathbb{P}(\xi_1 \neq 0, \xi_1 + \xi_2 \neq 0, \ldots, \xi_1 + \xi_2 + \cdots + \xi_j \neq 0)\\ &= \mathbb{P}(T_0 > j).\end{aligned}$$

#### A.3 Mean, variance and convergence in law of the range

We deal with the cases $d = 1, 2, 3$, considered separately, since very different behaviors occur.

##### A.3.1 Case $d = 1$

It is straightforward that

$$\mathtt{N}_n = 1 + \max_{0\leq k\leq n} X_k - \min_{0\leq k\leq n} X_k = 1 + \max\{X_p - X_q : 0 \leq p \leq n, 0 \leq q \leq n\}.$$

One expects that $\mathbb{E}(\mathtt{N}_n) = \mathcal{O}(\sqrt{n}\,)$. In [3] the authors obtain the following *exact* formula:

$$\mathbb{E}(\mathtt{N}_n) = \frac{\binom{2\lfloor\frac{n}{2}\rfloor}{\lfloor\frac{n}{2}\rfloor}}{2^{2\lfloor\frac{n}{2}\rfloor}} \left(n + 2\left\lfloor\tfrac{n}{2}\right\rfloor + 1\right),\; n \geq 1.$$

Stirling approximation $n! \underset{n\to+\infty}{\sim} n^n\, \mathrm{e}^{-n} \sqrt{2\pi n}$ yields

$$\mathbb{E}(\mathtt{N}_n) \underset{n\to+\infty}{\sim} \sqrt{\frac{8}{\pi}}\,\sqrt{n}. \tag{A.7}$$

For the variance we have (see [4])

$$\mathrm{Var}(\mathtt{N}_n) \underset{n\to+\infty}{\sim} 4\left(\log 2 - \frac{2}{\pi}\right) n. \tag{A.8}$$

The random variable $\mathtt{N}_n / \sqrt{n}$ converges in law to a non-degenerate probability distribution ([5],see also [4]), namely, we have the following result.

**Theorem A.6**. *We have*

$$\frac{\mathtt{N}_n}{\sqrt{n}} \xrightarrow{\text{law}} \mathcal{F} := \max_{0\leq s\leq 1} \mathcal{B}_s^{(1)} - \min_{0\leq s\leq 1} \mathcal{B}_s^{(1)}, \tag{A.9}$$

*where* $(\mathcal{B}_s^{(1)})_{s\geq 0}$ *is the standard one-dimensional Brownian motion.*

Compare this convergence in law with Theorem A.3 in view of (A.7). We see that $\mathtt{N}_n / \mathbb{E}(\mathtt{N}_n)$ doesn't converge to $1$ with probability one, but converges in law to a non-degenerate random variable. There is a formula for the probability density function $\rho$ of the random variable $\mathcal{F}$, namely

$$\rho(u) = \frac{8}{\sqrt{2\pi}} \sum_{m=1}^{+\infty} (-1)^{m-1} m^2\, \mathrm{e}^{-\frac{m^2 u^2}{2}}, u \in \mathbb{R}_+\,. \tag{A.10}$$

(So the cumulative distribution function of $\mathcal{F}$ is continuous.) This is proved by Feller [4].

We end this section by an important remark.

**Remark A.7.** *Looking at the proof of Theorem 6.2 in [5], we have that* $\mathbb{E}(\mathtt{N}_n/\sqrt{n}) \xrightarrow[n\to+\infty]{} \mathbb{E}(\mathcal{F}) = \sqrt{8/\pi}$ *and* $\mathrm{Var}(\mathtt{N}_n/\sqrt{n}) \xrightarrow[n\to+\infty]{} \mathrm{Var}(\mathcal{F}) = 4(\log 2 - 2/\pi)$. *This follows from a uniform bound in* $n$ *of the third moment of* $\mathtt{N}_n/\sqrt{n}$, *which implies that the square of* $\mathtt{N}_n/\sqrt{n}$ *is uniformly integrable. This is a well-known sufficient condition to ensure, from the convergence in law, the convergence of the first two moments of* $\mathtt{N}_n/\sqrt{n}$ *toward the first two moments of* $\mathcal{F}$. *Another way to proceed is to compute directly the first and second moments of* $\mathcal{F}$ *using its density, and to see that the results match with* (A.7) *and* (A.8). *Note also that, by classical properties of the Brownian motion, one has* $\mathbb{E}(\mathcal{F}) = 2\mathbb{E}(|\mathcal{B}_1^{(1)}|) = 2\sqrt{2/\pi}$ *(twice the expected value of the standard half-normal distribution).*

##### A.3.2 Case $d = 2$

We first recall the classical approximation [2]:

$$\mathbb{E}(\mathtt{N}_n) = \frac{\pi n}{\log n}\left(1 + \mathcal{O}\left(\frac{\log\log n}{\log n}\right)\right). \tag{A.11}$$

In fact, in order to obtain precise enough approximation of the number of prey consumed (see Section B.2 below) we need an approximation at order 2 for $\mathbb{E}(\mathtt{N}_n)$ that we couldn't find in the literature, so we provide a proof.

**Proposition A.8.** *For all* $n \geq 2$, *we have*

$$\mathbb{E}(\mathtt{N}_n) = \frac{\pi n}{\log n}\left(1 - \frac{\gamma}{\log n} + \mathcal{O}\left(\frac{1}{(\log n)^2}\right)\right) \tag{A.12}$$

*where* $\gamma$ *is the Euler-Mascheroni constant* ($\gamma = -\int_0^\infty \log(u)\,\mathrm{e}^{-u}\,\mathrm{d}u \approx 0,577$).[2]

*Proof.* We use [6, Theorem 1.3] which gives for any $k \in 2\mathbb{N}$

$$\mathbb{P}(T_0 = k) = \frac{\pi}{k(\log k)^2}\left(1 - \frac{2\gamma}{\log k} + \mathcal{O}\left(\frac{1}{(\log k)^2}\right)\right). \tag{A.13}$$

Hence for all $j \geq 2$ we have

$$\mathbb{P}(T_0 > j) = \sum_{k=j+1}^{+\infty} \mathbb{P}(T_0 = k) = \frac{\pi}{\log j}\left(1 - \frac{\gamma}{\log j} + \mathcal{O}\left(\frac{1}{(\log j)^2}\right)\right)$$

where we approximate in the standard way the series by an integral. Using (A.6) yields (A.12) after some standard but tedious calculations. More precisely, we need an asymptotic expansion for, *e.g.*,

$$\sum_{j=2}^{n} \frac{1}{\log j}$$

(which goes to $+\infty$ when $n \to +\infty$.) The basic idea is to approximate this sum by an integral, and then integrate by parts. Then one can use the following expansion for the offset logarithmic integral:

$$\mathrm{Li}(x) := \int_2^x \frac{\mathrm{d}u}{\log u} = \frac{x}{\log x} + \frac{1!\,x}{(\log x)^2} + \cdots + \frac{(k-1)!\,x}{(\log x)^k} + \mathcal{O}\left(\frac{x}{(\log x)^{k+1}}\right).$$

[2] Note that log stands for the natural logarithm.

We refer to [7, Section 3.3] for a nice exposition. □

Concerning the variance, Jain and Pruitt [5] proved that

$$\mathrm{Var}(\mathtt{N}_n) \underset{n\to+\infty}{\sim} \frac{2\pi^2 K\, n^2}{(\log n)^4} \tag{A.14}$$

where

$$K = -\int_0^1 \frac{\log u}{1-u+u^2}\,\mathrm{d}u + \frac{1}{2} - \frac{\pi^2}{12} \approx 0.8495. \tag{A.15}$$

Finally, Le Gall [8] proved the following result. We introduce the short-hand notation

$$Z_n := \frac{(\log n)^2}{n}\big(\mathtt{N}_n - \mathbb{E}(\mathtt{N}_n)\big). \tag{A.16}$$

**Theorem A.9.** *We have*

$$Z_n \xrightarrow{\text{law}} -2\pi^2 \mathcal{L}_{\mathcal{G}} \tag{A.17}$$

*where the random variable $\mathcal{L}_{\mathcal{G}}$ is the "renormalized self-intersection local time" for planar Brownian motion.*

Formally,

$$\mathcal{L}_{\mathcal{G}} = \iint_{\{0\le s<t\le 1\}} \delta_{(0)}(\mathcal{B}_s^{(2)} - \mathcal{B}_t^{(2)})\,\mathrm{d}s\,\mathrm{d}t - \mathbb{E}\left(\iint_{\{0\le s<t\le 1\}} \delta_{(0)}(\mathcal{B}_s^{(2)} - \mathcal{B}_t^{(2)})\,\mathrm{d}s\,\mathrm{d}t\right)$$

where $(\mathcal{B}_t^{(2)})_{t\ge 0}$ is a two-dimensional (standard) Brownian motion. Also formally $\mathbb{E}(\mathcal{L}_{\mathcal{G}}) = 0$, as it should be since it is the limit of centered random variables. Notice the unusual normalization by $n/(\log n)^2$, and the fact that the limiting random variable is not Gaussian. An elementary rigorous construction of $\mathcal{L}_{\mathcal{G}}$ is provided in [9].

The following uniform control of the exponential moment of the random variables $Z_n$ will be essential below. See [10] (Theorem 5.4 p. 26) for a proof.

**Theorem A.10.** *There exists $\theta > 0$ such that*

$$\sup_{n\in\mathbb{N}} \mathbb{E}\left(\exp\left(\theta|Z_n|\right)\right) < +\infty. \tag{A.18}$$

By standard arguments, we deduce the following corollaries.

**Corollary A.1.** *There exist $c_1, c_2 > 0$ such that*

$$\sup_{n\in\mathbb{N}} \mathbb{P}\big(|Z_n| > u\big) \le c_1\,\mathrm{e}^{-c_2 u},\ u > 0. \tag{A.19}$$

*Proof.* By Markov inequality we have (since $\theta > 0$)

$$\mathbb{P}\big(|Z_n| > u\big) = \mathbb{P}\big(\theta|Z_n| > \theta u\big) = \mathbb{P}\left(\mathrm{e}^{\theta|Z_n|} > \mathrm{e}^{\theta u}\right) \le \mathbb{E}\left(\exp\left(\theta|Z_n|\right)\right)\mathrm{e}^{-\theta u}.$$

This proves (A.19) with $c_1 = \sup_{n\in\mathbb{N}} \mathbb{E}\left(\exp\left(\theta|Z_n|\right)\right)$ and $c_2 = \theta$. □

**Corollary A.2.** *We have*

$$\lim_{n\to+\infty} \mathbb{E}\big(Z_n^2\big) = \mathbb{E}\big((-2\pi^2\mathcal{L}_{\mathcal{G}})^2\big) = \mathrm{Var}(2\pi^2\mathcal{L}_{\mathcal{G}}) = 2\pi^2 K \tag{A.20}$$

*where $K$ is defined in* (A.15).

*Proof.* A classical condition implying that convergence in law of $Z_n$ to $-2\pi^2\mathcal{L}_{\mathcal{G}}$ enforces convergence of $\mathbb{E}(Z_n^2)$ to $\mathbb{E}\big((-2\pi^2\mathcal{L}_{\mathcal{G}})^2\big)$ is uniform integrability of $\big(Z_n^2\big)_{n\in\mathbb{N}}$ (see Theorem 3.5 p. 31 in [11]). Using Corollary A.1, it is straightfoward to obtain $\sup_{n\in\mathbb{N}} \mathbb{E}\big(Z_n^3\big) < +\infty$, which implies the desired uniform integrability. □

**Remark A.11.** *Notice that the previous proof actually shows that, for all $p \geq 1$, $\mathbb{E}(|Z_n|^p)$ converges to the $p$-th moment of $-2\pi^2\mathcal{L}_{\mathcal{G}}$.*

##### A.3.3 Case $d = 3$

Dvoretsky and Erdös [2] proved that

$$\mathbb{E}(\mathtt{N}_n) = \kappa(3)\, n + \mathcal{O}\big(\sqrt{n}\,\big), \tag{A.21}$$

where $\kappa(3) \approx 0.6595$ (recall that this is the probability that the walk never returns to the origin). This is consistent with (A.2). Jain and Pruitt [12] (see Theorems 2 and 4) proved the following results.

**Theorem A.12.** *We have*

$$\frac{\mathtt{N}_n - \kappa(3)n}{\sigma\sqrt{n\log n}} \xrightarrow{\text{law}} \mathcal{N}(0,1) \tag{A.22}$$

*where $\mathcal{N}(0,1)$ is a standard Gaussian random variable (mean $0$ and variance $1$), and*

$$\sigma = \frac{3\sqrt{3}\,\kappa(3)^2}{\sqrt{2}\pi}. \tag{A.23}$$

*We also have*

$$\mathrm{Var}(\mathtt{N}_n) \underset{n\to+\infty}{\sim} \sigma^2\, n\log n.$$

#### A.4 More general random walks

For the sake of simplicity, we considered only the simplest random walks, namely, simple symmetric random walks. In fact, all the previous results hold true for a much more general class of random walks, up to minor changes, that we briefly describe. We refer to the book of [13] for more informations on random walks. We consider increments $\xi_k$'s ($k \geq 1$) that are independent and identically distributed random variables taking values in $\mathbb{Z}^d$, not necessarily in $\{\pm\, \mathrm{e}^{(1)}, \ldots, \pm\, \mathrm{e}^{(d)}\}$. (The case treated so far is the very special case when, if the walk is at a given site, the next move is to choose one of the $2d$ nearest neighbors of that site with probability $(2d)^{-1}$.) The corresponding random walk is defined by $X_0 = 0$ (so we suppose that the walk starts at the origin of $\mathbb{Z}^d$), and $X_n = \sum_{k=1}^n \xi_k$, $n = 1, 2, \ldots$. We assume that $\mathbb{E}(\xi_1) = 0$ and $\mathbb{E}(|\xi_1|^2) < +\infty$.[3]

[3] For $i \in \mathbb{Z}^d$, $|i|^2 := (i^{(1)})^2 + \cdots + (i^{(d)})^2$.

An important subclass of random walks with these two properties is the class of finite-range symmetric random walks. This means that there exists a finite set $V = \{i_1, \dots, i_r\} \subset \mathbb{Z}^d \backslash \{0\}$ which is generating (that is, every $j \in \mathbb{Z}^d$ can be written as $\alpha_1 i_1 + \cdots + \alpha_r i_r$ for some $\alpha_1, \dots, \alpha_r \in \mathbb{Z}$, so that the walk will be irreducible, and a function $\varrho : V \to ]0,1]$ with $\varrho(i_1) + \cdots + \varrho(x_r) = 1$, defining a symmetric probability distribution $p$ on $\mathbb{Z}^d$ as follows: $p(i_k) = p(-i_k) = \frac{1}{2}\varrho(i_k)$ $(i_k \in V)$. (Notice that we consider only the case $p(0) = 0$, that is, the walk has to leave the site where it is with probability one.) Given $p$, one can define $(X_n)_{n\geq 0}$ as the (time-homogeneous) Markov chain with state space $\mathbb{Z}^d$ and transition probabilities $\mathbb{P}(X_{n+1} = \ell | X_n = k) = p(\ell - k)$, or as $X_n = \sum_{k=1}^n \xi_k$ where the $\xi_k$'s are independent random variables with distribution $p$. The simple symmetric random walk is the case where $V = \{\mathrm{e}^{(1)}, \dots, \mathrm{e}^{(d)}\}$.

We now indicate how the results of Section A.3 generalize.

We denote by $Q = \big(\mathbb{E}(\xi_1^{(i)} \xi_1^{(j)})\big)_{1\leq i,j \leq d}$ the covariance matrix, and we let $\sigma^2 := \big(\det Q\big)^{1/d}$.

Theorem A.2 remains true for the class of random walks described above. In fact, having integrable increments is enough, see [1, p. 38].

Theorem A.3 can be generalized in dimension $d = 2$ for any random walk as above, provided it is recurrent, see Theorem 3.1 in [5].

Concerning Section A.3.1 ($d = 1$), Theorem A.6 is true for any random walk as defined above, up to replacing the interval $0 \leq s \leq 1$ by $0 \leq s \leq \sigma^2$.

Concerning the results listed in Section A.3.2 ($d = 2$), one has $\lim_{n\to+\infty} (\log n/n)\ \mathtt{N}_n = 2\pi\sigma^2$ almost-surely. Approximation (A.12) relies on (A.13) which remains the same, up to a multiplicative constant:

$$\mathbb{P}(T_0 = k) = \frac{2\pi\sigma^2}{k(\log k)^2} \left(1 - \frac{2\gamma}{\log k} + \mathcal{O}\left(\frac{1}{(\log k)^2}\right)\right).$$

This is because we can use [6, Theorem 1.3].

Theorem A.9 remains true with $-2\pi^2 \mathcal{L}_{\mathcal{G}}$ replaced by $-4\pi^2\sigma^2 \mathcal{L}_{\mathcal{G}}$, see [8]. Regarding the variance, (A.14) remains the same, up to the constant (see [5], Theorem 4.2): $\mathrm{Var}(\mathtt{N}_n) \underset{n\to+\infty}{\sim} \frac{c\,n^2}{(\log n)^4}$ where $c = 8\pi^2 K \sigma^4$ (where $K$ is defined in (A.15)).

Concerning Section A.3.3 ($d = 3$), we get the same behavior for the above class of random walks, up to constants that depend on $\xi_1$, see [14].

**Remark A.13** (Random walks with a drift). *The situation is very different when $\mathbb{E}(\xi_1) \neq 0$ (non-zero drift). In that case, the walk is strongly transient regardless of the dimension. We recall that a random walk is either transient or recurrent. Transience means that the probability that the walk never comes back to the origin is strictly positive, that is, $\kappa(d) > 0$ (see (A.1)). There are recurrent random walks only if the dimension is one or two. The simple symmetric random walk in dimensions 1 and 2 is recurrent. We will not define strong transience here. For instance, every random walk such that $\mathbb{E}(\xi_1) \neq 0$ and $\mathbb{E}(|\xi_1|^2) < +\infty$ is strongly transient. We just want to emphasize that, in that case, discarding the trivial case where the range grows deterministically (which is when $\kappa(d) = 1$), its variance grows like a constant times $n$, and a* *central limit theorem in the classical form is true: there exists* $\sigma^2 > 0$ *such that* $(\mathtt{N}_n - n\kappa(d))/(\sigma\sqrt{n}) \xrightarrow{\text{law}} N(0,1)$, *see [14] for details.*

**Remark A.14**. *Notice that the random walk may take place on a proper subgroup of* $\mathbb{Z}^d$. *In this case, the subgroup is isomorphic to some* $\mathbb{Z}^{d'}$ *for* $d' \le d$. *If* $d' < d$, *then the transformation should be made and the problem considered in* $d'$ *dimensions. We implicitly assumed that this reduction had been made, if necessary, and that* $d$ *is the genuine dimension of the random walk. (This means in particular that the determinant of the covariance matrix of the random walk is not equal to* $0$.*)*

### B The number of consumed prey

All proofs and results given in this section are new.

#### B.1 Process dual to the range

Observing that $(\mathtt{N}_n)_{n\ge 1}$ is an increasing process, it is natural to define its "inverse", namely

$$S_k = \inf\{n \ge k : \mathtt{N}_n \ge k\},\ k \ge 1. \tag{B.1}$$

This is the smallest number of steps for the walk to visit at least $k$ distinct sites. We have the following fundamental "duality relation"

$$\{\mathtt{N}_n \le k\} = \{n \le S_k\}. \tag{B.2}$$

In particular, studying the law of $\mathtt{N}_n$ is equivalent to studying the law of $S_k$. Observe that, for each $k$, $S_k$ is a stopping time.

#### B.2 Connecting the number of consumed prey to the range of the random walk

We define two important parameters:

- $\tau_e$, defined as the time needed for a predator to go across an edge of the lattice $\mathbb{Z}^d$
- $\tau_h$, defined as the handling time, *i.e.*, the time taken by the predator to consume one prey.

The dimensionless parameter $\tau_h/\tau_e$ will show up in many instances. We will always assume that $\tau_e > 0$ ($\tau_h$ can be equal to $0$). The first one gives a time scale with respect to which we can say what a large time $t$ means. Then, we define the stochastic process giving the evolution of our system. It is the joint process made of the random walk $(X_k)_{k\ge 0}$ and the corresponding sequence of (random) times $(T_k)_{k\ge 0}$ at which the walk (the predator) visits the sites.

Let $(X_k, T_k)_{k\geq 0}$ be the double sequence of random variables giving the sites that the predator visits (that is, the position of the random walk), together with the corresponding arrival times. The random walk $(X_k)_{k\geq 0}$ we consider is the simple symmetric random walk defined in the previous section (recall that $X_0 = (0, \ldots, 0)$). We can recursively define the arrival times by setting $T_0 = 0$, and letting

$$T_{k+1} = \begin{cases} T_k + \tau_h + \tau_e & \text{if} \quad X_k \notin \{X_0, X_1, \ldots X_{k-1}\} \\ T_k + \tau_e & \text{otherwise.} \end{cases}$$

The condition $X_k \notin \{X_0, X_1, \ldots X_{k-1}\}$ is equivalent to $\mathtt{N}_k > \mathtt{N}_{k-1}$. Then, the number of prey found before time $t$ is

$$R_t = \text{Card}\big\{k \geq 0 : T_k \leq t,\, \mathtt{N}_k > \mathtt{N}_{k-1}\big\} = \text{Card}\big\{k \geq 0 : T_k \leq t, X_k \notin \{X_1, \ldots X_{k-1}\}\big\}.$$

The functional response (the consumption rate) is then defined as

$$F_t = \frac{R_t}{t}.$$

Now, by definition of $R_t$ and using (B.2), we have

$$\{R_t > k\} = \{S_k \tau_e + (k-1)\,\tau_h < t\} = \left\{S_k < \frac{t}{\tau_e} - (k-1)\frac{\tau_h}{\tau_e}\right\} = \big\{\mathtt{N}_{n(t)} > k\big\} \tag{B.3}$$

where

$$n(t) := n_k(t) = \left\lfloor \frac{t}{\tau_e} - (k-1)\frac{\tau_h}{\tau_e} \right\rfloor \tag{B.4}$$

where $\lfloor \cdot \rfloor$ denotes integer part. (For $n(t)$ to be larger than or equal to 1, there is an obvious constraint on $t$ and $k$, namely $\frac{t}{\tau_e} \geq (k-1)\frac{\tau_h}{\tau_e} + 1$.)

##### B.2.1 Case $d = 1$

We have the following result.

**Theorem B.1.** *As $t \to +\infty$, we have*

$$\frac{R_t}{\sqrt{t/\tau_e}} \xrightarrow{\text{law}} \max_{0\leq s\leq 1} \mathcal{B}_s^{(1)} - \min_{0\leq s\leq 1} \mathcal{B}_s^{(1)} \tag{B.5}$$

*where $(\mathcal{B}_s^{(1)})_{s\geq 0}$ is the standard one-dimensional Brownian motion.*

Observe that $\tau_h$ plays no role in the limiting random variable.

*Proof.* Let $t > 0$. For convenience, we set $t_e := t/\tau_e$ (dimensionless parameter counting time in units of $\tau_e$). Let $u \in \mathbb{R}$. Using (B.3) and (B.4), we have

$$\mathbb{P}\left(\frac{R_t}{\sqrt{t_e}} \leq u\right) = \mathbb{P}\left(\sqrt{\frac{n(t)}{t_e}}\, \frac{\mathtt{N}_{n(t)}}{\sqrt{n(t)}} \leq u\right) \tag{B.6}$$

where

$$n(t) = \left\lfloor t_e - \left(u\sqrt{t_e} - 1\right) \frac{\tau_h}{\tau_e} \right\rfloor.$$

Since $\sqrt{\frac{n(t)}{t_e}} \to 1$ as $t \to +\infty$, using Slutsky's theorem and Theorem A.6, we obtain

$$\sqrt{\frac{n(t)}{t_e}} \frac{\mathtt{N}_{n(t)}}{\sqrt{n(t)}} \xrightarrow{\text{law}} \mathcal{F}$$

where $\mathcal{F} = \max_{0\le s\le 1} \mathcal{B}_s^{(1)} - \min_{0\le s\le 1} \mathcal{B}_s^{(1)}$. This means that the right-hand side in (B.6) goes to $\mathbb{P}(\mathcal{F} \le u)$ (its cumulative distribution function is continuous since $\mathcal{F}$ has a density). [4] But, since $u \in \mathbb{R}$ is arbitrary, we obtain that the left-hand side in (B.6) goes to $\mathbb{P}(\mathcal{F} \le u)$ for all $u \in \mathbb{R}$, which entails the convergence in law of $R_t/\sqrt{t_e}$ to $\mathcal{F}$. □

The first two moments of $R_t/\sqrt{t/\tau_e}$ do converge to those of $\max_{0\le s\le 1} \mathcal{B}_s^{(1)} - \min_{0\le s\le 1} \mathcal{B}_s^{(1)}$.

**Proposition B.2.** *We have*

$$\mathbb{E}\left(\frac{R_t}{\sqrt{t/\tau_e}}\right) \xrightarrow[t\to+\infty]{} \mathbb{E}\left(\max_{0\le s\le 1} \mathcal{B}_s^{(1)} - \min_{0\le s\le 1} \mathcal{B}_s^{(1)}\right) = 2\sqrt{\frac{2}{\pi}} \approx 1.5958,$$

*and*

$$\operatorname{Var}\left(\frac{R_t}{\sqrt{t/\tau_e}}\right) \xrightarrow[t\to+\infty]{} \operatorname{Var}\left(\max_{0\le s\le 1} \mathcal{B}_s^{(1)} - \min_{0\le s\le 1} \mathcal{B}_s^{(1)}\right) = 4\left(\log 2 - \frac{2}{\pi}\right) \approx 0.2261.$$

*Proof.* In view of Remark A.7, we show how to control the third moment of $\widetilde{R}_t := R_t/\sqrt{t_e}$ (where $t_e = t/\tau_e$), uniformly in $t$. Since $\widetilde{R}_t$ is a positive random variable, we only need to control the tail probability $\mathbb{P}\big(\widetilde{R}_t > u\big)$ for $u > 0$. Taking $n(t) = \left\lfloor t_e - (k-1)\frac{\tau_h}{\tau_e} \right\rfloor$ with $k = k(t,u) = u\sqrt{t_e}$, we get at once using (B.3) and the obvious inequality $\mathtt{N}_{n(t)} \le \mathtt{N}_{\lfloor t_e \rfloor}$ that

$$\mathbb{P}\big(\widetilde{R}_t > u\big) = \mathbb{P}\big(R_t > k\big) \le \mathbb{P}\big(\mathtt{N}_{\lfloor t_e \rfloor} > k\big) = \mathbb{P}\left(\frac{\mathtt{N}_{\lfloor t_e \rfloor}}{\sqrt{t_e}} > u\right).$$

We conclude by using Theorem 6.2 in [5] which states that $\mathbb{E}\left[\left(\frac{\mathtt{N}_{\lfloor t_e \rfloor}}{\sqrt{t_e}}\right)^3\right] = \mathcal{O}(1)$, and the formula $\mathbb{E}\big(\widetilde{R}_t^3\big) = 2\int_0^{+\infty} u^2 \mathbb{P}\big(\widetilde{R}_t > u\big)\,\mathrm{d}u$. □

##### B.2.2 Case $d = 2$

The situation for $d = 2$ is completely different from that in dimension $d = 1$.

**Theorem B.3.** *We have*

$$\frac{\left(\log\left(\frac{t}{\tau_e}\right)\right)^2}{\frac{t}{\tau_e}} \left(R_t - \frac{\pi \frac{t}{\tau_e}}{\log\left(\frac{t}{\tau_e}\right)}\right) \xrightarrow{\text{law}} -\pi\left(2\pi\mathcal{L}_{\mathcal{G}} + \frac{\pi\,\tau_h}{\tau_e} + \gamma\right), \text{ as } t \to +\infty \tag{B.7}$$

*where $\mathcal{L}_{\mathcal{G}}$ is the random variable appearing in Theorem A.9.*

[4] Recall that if $(Y_n)_{n\ge 0}$ is a sequence of real-valued random variables, and $Y$ a real-valued random variable, then $Y_n \xrightarrow{\text{law}} Y$ if and only if $\mathbb{P}(Y_n \le u) \to \mathbb{P}(Y \le u)$, as $n \to +\infty$, at every point where $u \mapsto \mathbb{P}(Y \le u)$ is continuous.

*Proof.* Let $t > 0$. As before, we set $t_e := t/\tau_e$. To alleviate notation, let $a(t) := (\log t)^2/t$, and

$$\widetilde{R}_t := a(t_e)\left(R_t - \frac{\pi t_e}{\log t_e}\right).$$

Using (A.12), we rewrite the left-hand side of (B.7):

$$\widetilde{R}_t = a(t_e)\left(R_t - \mathbb{E}\left(\mathtt{N}_{\lfloor t_e \rfloor}\right)\right) - \pi\gamma + \mathcal{O}\left(\frac{1}{\log t_e}\right). \tag{B.8}$$

Hence, by Slutsky's theorem, if $a(t_e)\left(R_t - \mathbb{E}\left(\mathtt{N}_{\lfloor t_e \rfloor}\right)\right)$ converges in law to a random variable, say, $\mathcal{W}$, then $\widetilde{R}_t$ converges in law to

$$\mathcal{W} - \pi\gamma. \tag{B.9}$$

We now want to use (B.3), that is

$$\{R_t \le k\} = \{\mathtt{N}_{n(t)} \le k\}, \tag{B.10}$$

where we take

$$n(t) = n(t,u) = \left\lfloor t_e - (k-1)\frac{\tau_h}{\tau_e} \right\rfloor \quad \text{and} \quad k = k(t,u) := \left\lfloor a(t_e)^{-1}u + \mathbb{E}\left(\mathtt{N}_{\lfloor t_e \rfloor}\right)\right\rfloor,$$

and where $u \in \mathbb{R}$ is a continuity point of the cumulative distribution function of $-2\pi^2\mathcal{L}_{\mathcal{G}}$. Hence, the equality between the events in (B.10) writes

$$\left\{a(t_e)\left(R_t - \mathbb{E}\left(\mathtt{N}_{\lfloor t_e \rfloor}\right)\right) \le u\right\} = \left\{a(t_e)\left(\mathtt{N}_{n(t)} - \mathbb{E}\left(\mathtt{N}_{\lfloor t_e \rfloor}\right)\right) \le u\right\}. \tag{B.11}$$

Using Theorem A.9, with $n$ replaced by $n(t)$, we have

$$a(n(t))\left(\mathtt{N}_{n(t)} - \mathbb{E}(\mathtt{N}_{n(t)})\right) \xrightarrow{\text{law}} -2\pi^2\mathcal{L}_{\mathcal{G}}. \tag{B.12}$$

(Observe that $n(t) \to +\infty$, as $t \to +\infty$, since $\mathbb{E}\left(\mathtt{N}_{\lfloor t_e \rfloor}\right) \underset{t\to+\infty}{\sim} (\pi t_e)/\log t_e$ by (A.11), whence $n(t) \underset{t\to+\infty}{\sim} t_e$.) To use (B.12), we write

$$a(t_e)\left(\mathtt{N}_{n(t)} - \mathbb{E}\left(\mathtt{N}_{\lfloor t_e \rfloor}\right)\right) = \frac{a(t_e)}{a(n(t))}\left(a(n(t))\left(\mathtt{N}_{n(t)} - \mathbb{E}\left(\mathtt{N}_{n(t)}\right)\right) + a(t_e)\,\mathbb{E}\left(\mathtt{N}_{n(t)} - \mathtt{N}_{\lfloor t_e \rfloor}\right)\right). \tag{B.13}$$

Using (A.11), and the fact that $k \underset{t\to+\infty}{\sim} \pi\, t_e/\log(t_e)$, we have

$$\mathbb{E}\left(\mathtt{N}_{\lfloor t_e \rfloor}\right) - \mathbb{E}\left(\mathtt{N}_{n(t)}\right) \underset{t\to+\infty}{\sim} \frac{\pi}{\log t_e}\frac{\tau_h k}{\tau_e} \underset{t\to+\infty}{\sim} \frac{\tau_h t}{\tau_e^2}\frac{\pi^2}{(\log t_e)^2}.$$

Hence

$$a(t_e)\,\mathbb{E}\left(\mathtt{N}_{n(t)} - \mathtt{N}_{\lfloor t_e \rfloor}\right) \underset{t\to+\infty}{\sim} -\pi^2\frac{\tau_h}{\tau_e}. \tag{B.14}$$

We leave to the reader check that $a(n(t))/a(t_e) \underset{t\to+\infty}{\sim} 1$. Then, we apply Slutsky's theorem once more, together with (B.12), to obtain that the right-hand side of (B.13) converges in law to $-2\pi^2\mathcal{L}_{\mathcal{G}} - (\pi^2\tau_h)/\tau_e$. Hence we obtain by (B.13) that

$$a(t_e)\left(\mathtt{N}_{n(t)} - \mathbb{E}\left(\mathtt{N}_{\lfloor t_e \rfloor}\right)\right) \xrightarrow{\text{law}} -2\pi^2\mathcal{L}_{\mathcal{G}} - \pi^2\frac{\tau_h}{\tau_e}.$$

It follows that $\mathbb{P}\big(a(t_e)\left((\mathtt{N}_{n(t)} - \mathbb{E}\left(\mathtt{N}_{\lfloor t_e \rfloor}\right)\right) \le u\big) \xrightarrow[t\to+\infty]{} \mathbb{P}\big(-2\pi^2\mathcal{L}_{\mathcal{G}} - (\pi^2\tau_h)/\tau_e \le u\big)$ for any $u \in \mathbb{R}$ is an arbitrary continuity point of the cumulative distribution function. We deduce, using (B.11), that $a(t_e)\big(R_t - \mathbb{E}\left(\mathtt{N}_{\lfloor t_e \rfloor}\right)\big) \xrightarrow{\text{law}} -2\pi^2\mathcal{L}_{\mathcal{G}} - (\pi^2\tau_h)/\tau_e$. Therefore, coming back to (B.8) (see the sentence right after it, and also (B.9)), we finally obtain that $\mathcal{W} = -2\pi^2\mathcal{L}_{\mathcal{G}} - (\pi^2\tau_h)/\tau_e$, which entails that

$$\widetilde{R}_t \xrightarrow{\text{law}} -2\pi^2\mathcal{L}_{\mathcal{G}} - \frac{\pi^2\tau_h}{\tau_e} - \pi\gamma$$

which after rearrangement is exactly (B.7), which ends the proof. □

The following result is the analog of Corollary A.2, and its proof goes along the same lines.

**Proposition B.4.** *We have*

$$\mathbb{E}(R_t) \underset{t\to+\infty}{\sim} \frac{\pi t_e}{\log t_e} \tag{B.15}$$

*and*

$$\left(\frac{\left(\log\left(\frac{t}{\tau_e}\right)\right)^2}{\frac{t}{\tau_e}}\right)^2 \operatorname{Var}(R_t) \xrightarrow[t\to+\infty]{} 2\pi^2 K, \tag{B.16}$$

*where $K$ is defined in* (A.15).

*Proof.* As before, we use the notations $t_e = t/\tau_e$, $a(t) = (\log t)^2/t$, and

$$\widetilde{R}_t := a(t_e)\left(R_t - \frac{\pi t_e}{\log t_e}\right).$$

We first show how to control $\mathbb{P}\big(\widetilde{R}_t > u\big)$ for all $u > 0$, uniformly in $t$ (which is the crucial point). We use again (B.3), that is,

$$\{R_t > k\} = \{\mathtt{N}_{n(t)} > k\}, \tag{B.17}$$

where we now take

$$n(t) = n(t,u) = \left\lfloor t_e - (k-1)\frac{\tau_h}{\tau_e}\right\rfloor, \quad k = k(t,u) := \left\lfloor a(t_e)^{-1}u + \frac{\pi t_e}{\log t_e}\right\rfloor.$$

Since $\mathtt{N}_{n(t)} \le \mathtt{N}_{\lfloor t_e \rfloor}$ we have

$$\mathbb{P}(R_t > k) = \mathbb{P}(\mathtt{N}_{n(t)} > k) \le \mathbb{P}(\mathtt{N}_{\lfloor t_e \rfloor} > k). \tag{B.18}$$

We now bound the rightmost term:

$$\begin{aligned}\mathbb{P}(\mathtt{N}_{\lfloor t_e \rfloor} > k) &= \mathbb{P}\left(a(t_e)\left(\mathtt{N}_{\lfloor t_e \rfloor} - \frac{\pi t_e}{\log t_e}\right) > u\right)\\ &= \mathbb{P}\left(a(t_e)\left(\mathtt{N}_{\lfloor t_e \rfloor} - \mathbb{E}(\mathtt{N}_{\lfloor t_e \rfloor})\right) > u + a(t_e)\left(\frac{\pi t_e}{\log t_e} - \mathbb{E}(\mathtt{N}_{\lfloor t_e \rfloor})\right)\right).\end{aligned}$$

We use again (A.12) to get

$$a(t_e)\left(\frac{\pi t_e}{\log t_e} - \mathbb{E}(\mathtt{N}_{\lfloor t_e \rfloor})\right) = \pi\gamma + \mathcal{O}\left(\frac{1}{\log t_e}\right),$$

which is larger than $\pi\gamma/2$ for all $t$ larger than some $t_0$ (independent of $u$). It follows from (B.18) that

$$\mathbb{P}\big(\widetilde{R}_t > u\big) = \mathbb{P}(R_t > k) \le \mathbb{P}\left(a(t_e)\left(\mathtt{N}_{\lfloor t_e \rfloor} - \mathbb{E}(\mathtt{N}_{\lfloor t_e \rfloor})\right) > u + \frac{\pi\gamma}{2}\right), \ t \ge t_0,$$

therefore, using (A.19), we obtain

$$\sup_{t\ge t_0} \mathbb{P}\big(\widetilde{R}_t > u\big) \le c_1\,\mathrm{e}^{-c_2 u}\,. \tag{B.19}$$

We now turn to show how to control $\mathbb{P}\big(\widetilde{R}_t < -u\big)$ for all $u > 0$, uniformly in $t$. We have

$$\mathbb{P}\big(\widetilde{R}_t < -u\big) = \mathbb{P}\left(a(t_e)\left(R_t - \frac{\pi t_e}{\log t_e}\right) < -u\right) = \mathbb{P}\left(R_t < -\frac{u}{a(t_e)} + \frac{\pi t_e}{\log t_e}\right) = \mathbb{P}(\mathtt{N}_{n(t)} < k)$$

where we used (B.3) with $k = k(t,u) := -a(t_e)u + (\pi t_e)/\log t_e$, whence

$$n(t) \ge \left\lfloor t_e + \left(\frac{u}{a(t_e)} - \frac{\pi t_e}{\log t_e}\right)\frac{\tau_h}{\tau_e}\right\rfloor \ge \left\lfloor t_e\left(1 - \frac{\pi\tau_h}{\tau_e}\frac{1}{\log t_e}\right)\right\rfloor =: n'(t).$$

(Observe that the rightmost term does not depend on $u$.) It follows that $\mathtt{N}_{n(t)} \ge \mathtt{N}_{\lfloor t_e(1-\frac{\pi\tau_h}{\tau_e}\frac{1}{\log t_e})\rfloor}$, hence

$$\begin{aligned}
\mathbb{P}\big(\widetilde{R}_t < -u\big) &\le \mathbb{P}\left(\mathtt{N}_{n'(t)} < k\right) \\
&= \mathbb{P}\left(a(t_e)\left(\mathtt{N}_{n'(t)} - \frac{\pi t_e}{\log t_e}\right) < -u\right) \\
&= \mathbb{P}\left(a(t_e)\left(\mathtt{N}_{n'(t)} - \mathbb{E}\left(\mathtt{N}_{n'(t)}\right)\right) < -u + a(t_e)\left(\frac{\pi t_e}{\log t_e} - \mathbb{E}\left(\mathtt{N}_{n'(t)}\right)\right)\right).
\end{aligned}$$

Once more, we use(A.12), and it is left to the reader to check that

$$a(t_e)\left(\frac{\pi t_e}{\log t_e} - \mathbb{E}\left(\mathtt{N}_{n'(t)}\right)\right) = -\pi\left(\pi\frac{\tau_h}{\tau_e} - \gamma\right) + \mathcal{O}\left(\frac{1}{\log t_e}\right).$$

There exists $t_0'$ (independent of $u$) such that for all $t \ge t_0'$ the right-hand side is less than or equal to $-(\pi^2\tau_h)/\tau_e + (3\pi\gamma)/2$, hence, using (A.19), we obtain

$$\mathbb{P}\big(\widetilde{R}_t < -u\big) \le c_1\,\mathrm{e}^{\frac{3\pi\gamma c_2}{2}}\,\mathrm{e}^{-c_2 u},$$

therefore

$$\sup_{t\ge t_0'} \mathbb{P}\big(\widetilde{R}_t < -u\big) \le c_1\,\mathrm{e}^{\frac{3\pi\gamma c_2}{2}}\,\mathrm{e}^{-c_2 u}\,. \tag{B.20}$$

Combining (B.19) and (B.20), we thus proved that

$$\sup_{t\ge t_0\vee t_0'} \mathbb{P}\big(\big|\widetilde{R}_t\big| > u\big) \le 2c_1\,\mathrm{e}^{\frac{3\pi\gamma c_2}{2}}\,\mathrm{e}^{-c_2 u}\,.$$

Reasoning as in the proof of Corollary A.2, we deduce that the expected value of the left-hand side of (B.7) converges to the expected value of its right-hand side, that is,

$$a(t_e)\left(\mathbb{E}(R_t) - \frac{\pi t_e}{\log t_e}\right) \to -\pi\left(\pi\frac{\tau_h}{\tau_e} + \gamma\right), \text{ as } t \to +\infty. \tag{B.21}$$

(Recall that $\mathbb{E}(\mathcal{L}_{\mathcal{G}}) = 0$.) This implies (B.15). We also know that

$$\mathbb{E}\big(\widetilde{R}_t^2\big) \to \mathbb{E}\left[\left(2\pi^2\mathcal{L}_{\mathcal{G}} + \frac{\pi^2\,\tau_h}{\tau_e} + \pi\gamma\right)^2\right] = \mathrm{Var}(2\pi^2\mathcal{L}_{\mathcal{G}}) + \left(\frac{\pi^2\,\tau_h}{\tau_e} + \pi\gamma\right)^2, \text{ as } t \to +\infty. \tag{B.22}$$

Since

$$a(t_e)^2 \operatorname{Var}(R_t) = \mathbb{E}\big(\widetilde{R}_t^2\big) - \left(a(t_e)\left(\mathbb{E}(R_t) - \frac{\pi t_e}{\log t_e}\right)\right)^2 \text{ as } t \to +\infty,$$

(B.16) follows from (B.21) and (B.22). The proof of the proposition is now complete. □

##### B.2.3 Case $d = 3$

We now deal with the case $d = 3$ which is completely different from the cases $d = 1$ and $d = 2$.

**Theorem B.5**. *We have*

$$\frac{R_t - \left(\frac{\kappa(3)}{1+\kappa(3)\frac{\tau_h}{\tau_e}}\right)\frac{t}{\tau_e}}{\sqrt{\frac{t}{\tau_e}\log\frac{t}{\tau_e}}} \xrightarrow{\text{law}} \mathcal{N}\big(0,\eta^2\big), \text{ as } t \to +\infty, \tag{B.23}$$

*where* $\mathcal{N}\big(0,\eta^2\big)$ *is a centered Gaussian random variable with variance*

$$\eta^2 = \frac{\sigma^2}{\left(1+\kappa(3)\frac{\tau_h}{\tau_e}\right)^3}, \tag{B.24}$$

*where* $\sigma$ *is given in* (A.23).

Observe that, when $\tau_h \to 0$, the coefficient of $t/\tau_e$ in (B.23) goes to $\kappa(3)$, and $\eta$ goes $\sigma$. So, in the limit $\tau_h \to 0$, $R_t$ behaves in the same way, in law, as $\mathtt{N}_n$ if $n$ is replaced by $t/\tau_e$ (see (A.22).

*Proof.* Let $t > 0$. Fix $u \in \mathbb{R}$ and recall that $t_e = t/\tau_e$. In view of (B.3) and (B.4) and the normalisation we are interested in for $R_t$, we choose $k = \lfloor u\,\sigma\sqrt{t_e\log t_e} + c\,t_e\rfloor$, where $c$ is a constant which will be chosen later on, and get

$$\mathbb{P}\left(\frac{R_t - c\,t_e}{\sigma\sqrt{t_e\log t_e}} \le u\right) = \mathbb{P}\left(\frac{\mathtt{N}_{n(t)} - c\,t_e}{\sigma\sqrt{t_e\log t_e}} \le u\right). \tag{B.25}$$

Now we rewrite the random variable that appears in the right-hand side of the previous equation as follows:

$$\frac{\mathtt{N}_{n(t)} - c\,t_e}{\sigma\sqrt{t_e\log t_e}} = \sqrt{\frac{n(t)\log n(t)}{t_e\log t_e}}\,\frac{\mathtt{N}_{n(t)} - \kappa(3)\,n(t)}{\sigma\sqrt{n(t)\log n(t)}} + \frac{1}{\sigma}\,\frac{\kappa(3)\,n(t) - c\,t_e}{\sqrt{t_e\log t_e}}. \tag{B.26}$$

We have

$$\kappa(3)\,n(t) - c\,t_e = \left(\kappa(3)\left(1 - \frac{c\tau_h}{\tau_e}\right) - c\right)t_e - \kappa(3)\,\sigma\frac{\tau_h}{\tau_e}\,u\sqrt{t_e\log t_e} + \mathcal{O}(1).$$

If the coefficient of $t_e$ is not equal to 0, then $(\kappa(3)\,n(t) - c\,t_e)/\sqrt{t_e\log t_e}$ blows up as $t \to +\infty$. To get a finite contribution in the limit, we look for a value of $c$ making this coefficient equal to 0, which works if

$$c = \frac{\kappa(3)}{1+\kappa(3)\frac{\tau_h}{\tau_e}}. \tag{B.27}$$

It is easy to check that

$$n(t) \underset{t\to+\infty}{\sim} \left(1 - \frac{c\,\tau_h}{\tau_e}\right)t_e = \frac{t_e}{1+\kappa(3)\frac{\tau_h}{\tau_e}}.$$

Hence

$$\sqrt{((n(t)\log n(t))/(t_e\log t_e)} \underset{t\to+\infty}{\sim} \frac{1}{\sqrt{1+\kappa(3)\frac{\tau_h}{\tau_e}}}.$$

We now apply Slutsky's theorem, together with Theorem A.12, to get that the right-hand in (B.26) side converges in law to

$$\frac{\mathcal{N}(0,1)}{\sqrt{1+\kappa(3)\frac{\tau_h}{\tau_e}}} - \frac{\kappa(3)\tau_h}{\tau_e}u$$

where $\mathcal{N}(0,1)$ is a standard Gaussian random variable. Since the limiting random variable has a density, its cumulative distribution function is continuous, hence we finally obtain (remember (B.26)):

$$\mathbb{P}\left(\frac{\mathtt{N}_{n(t)} - c\,t_e}{\sigma\sqrt{t_e\log t_e}} \le u\right) \xrightarrow[t\to+\infty]{} \mathbb{P}\left(\frac{\mathcal{N}(0,1)}{\sqrt{1+\kappa(3)\frac{\tau_h}{\tau_e}}} - \frac{\kappa(3)\tau_h}{\tau_e}u \le u\right) = \mathbb{P}\left(\mathcal{N}' \le u\right)$$

where $\mathcal{N}'$ is a centered Gaussian random variable with variance $1/\big(1+\kappa(3)\frac{\tau_h}{\tau_e}\big)^3$. In view of (B.25), this means that

$$\mathbb{P}\left(\frac{R_t - c\,t_e}{\sigma\sqrt{t_e\log t_e}} \le u\right) \xrightarrow[t\to+\infty]{} \mathbb{P}\left(\mathcal{N}' \le u\right)$$

where $c$ is given in (B.27). Since this is true for an arbitrary $u\in\mathbb{R}$, we thus proved that

$$\frac{R_t - \left(\frac{\kappa(3)}{1+\kappa(3)\frac{\tau_h}{\tau_e}}\right)t_e}{\sqrt{t_e\log t_e}} \xrightarrow{\text{law}} \sigma\,\mathcal{N}'$$

which concludes the proof, since the random variable $\sigma\,G'$ is equal in law to a centered Gaussian random variable with variance $\sigma^2/\big(1+\kappa(3)\frac{\tau_h}{\tau_e}\big)^3$. □

We finish this section by the following proposition.

**Proposition B.6.** *We have*

$$\mathbb{E}(R_t) \underset{t\to+\infty}{\sim} \left(\frac{\kappa(3)}{1+\kappa(3)\frac{\tau_h}{\tau_e}}\right)\frac{t}{\tau_e} \quad \textit{and} \quad \mathrm{Var}\left(\frac{R_t}{\sqrt{\frac{t}{\tau_e}\log\frac{t}{\tau_e}}}\right) \xrightarrow[t\to+\infty]{} \eta^2,$$

*where* $\eta$ *is defined in* (B.24).

We leave the proof to the reader. It follows the same pattern as before, namely the strategy consists in proving some uniform integrability, which in turn results from the uniform control of the 4th-order moment of $\big(\mathtt{N}_n - \mathbb{E}(\mathtt{N}_n)\big)/\sqrt{n\log n}$ proved in [5](see Theorem 4).

#### B.3 Coefficients of variation of $R_t$

Let $\sigma(R_t) := \sqrt{\mathrm{Var}(R_t)}$, that is, $\sigma(R_t)$ is the standard deviation of $R_t$. We now compute the coefficient of variation defined as $\mathrm{CV}_t^{(d)} = \sigma(R_t)/\mathbb{E}(R_t)$ for $d = 1,2,3$.

In dimension one, we deduce from Proposition B.2

$$\mathrm{CV}_t^{(1)} \underset{t\to+\infty}{\sim} \frac{2\sqrt{\log 2 - \frac{2}{\pi}}}{2\sqrt{\frac{2}{\pi}}} \approx 0.3.$$

In dimension two, Proposition B.4 implies that

$$\mathrm{CV}_t^{(d2} \underset{t\to+\infty}{\sim} \frac{\sqrt{2K}}{\log\frac{t}{\tau_e} - \gamma - \pi\frac{\tau_h}{\tau_e}} \approx \frac{1.3}{\log\frac{t}{\tau_e}}.$$

Finally, in dimension 3, Proposition B.6 gives

$$\mathrm{CV}_t^{(3)} \underset{t\to+\infty}{\sim} \frac{3\sqrt{3}\,\kappa(3)}{\sqrt{2}\,\pi\sqrt{1+\kappa(3)\frac{\tau_h}{\tau_e}}}\sqrt{\frac{\log\frac{t}{\tau_e}}{\frac{t}{\tau_e}}} \approx \frac{0.8}{\sqrt{1+0.7\frac{\tau_h}{\tau_e}}}\sqrt{\frac{\log\frac{t}{\tau_e}}{\frac{t}{\tau_e}}}.$$

#### B.4 Adding prey at random on the lattice

The same approach allows to extend the results on the asymptotic behavior of the functional response to the case where we drop at each site (independently of one another) a prey with probability $p$. We can follow the same argument as above and now use (B.29) and (B.30) below.

More precisely, we assume that, at each site, we draw a prey at random with probability $p$, where $p \in (0,1]$ is fixed. The prey are drawn independently of one another. Let us count the number of prey $\mathtt{N}_n^{(p)}$ seen by the predator after $n$ steps, *i.e.*, the number of different sites where there is a prey. Observe that $\mathtt{N}_n^{(1)} = \mathtt{N}_n$. Moreover, the process $\big(\mathtt{N}_n^{(p)}\big)_{n\geq 0}$ can be constructed using $(\mathtt{N}_n)_{n\geq 1}$ as follows. Let $(B_i)_{i\in\mathbb{Z}^d}$ be independent Bernoulli random variables with parameter $p$. The event $B_i = 1$ corresponds to the presence of a prey at site $i$. We denote by $Y_k \in \mathbb{Z}^d$ the value of the $k$-th distinct site visited by the random walk $(X_n)_{n\geq 0}$, so that $(Y_1,\dots,Y_{\mathtt{N}_n})$ are the sites successively visited after $n$ step. The key observation is that the random variables $V_k = B_{Y_k}$ also form a sequence of independent identically distributed Bernoulli random variables with parameter $p$, which is independent of $(\mathtt{N}_n)_{n\geq 1}$. Moreover

$$\mathtt{N}_n^{(p)} = \sum_{k=1}^{\mathtt{N}_n} V_k. \tag{B.28}$$

First, we immediately deduce that

$$\mathbb{E}\big(\mathtt{N}_n^{(p)}\big) = p\,\mathbb{E}(\mathtt{N}_n). \tag{B.29}$$

Next, letting $a_n^{(d)} = \mathbb{E}(\mathtt{N}_n)$, recall that $\mathtt{N}_n / a_n^{(d)}$ converges to $1$ with probability one in dimensions $d = 2,3$ (see (A.3), (A.11) and (A.21)). We deduce that

$$\mathbb{P}\left(\frac{\mathtt{N}_n^{(p)}}{a_n^{(d)}} \xrightarrow[n\to+\infty]{} p\right) = 1.$$

Indeed,

$$\frac{\mathtt{N}_n^{(p)}}{a_n^{(d)}} = \frac{\mathtt{N}_n}{a_n^{(d)}}\left(\frac{1}{\mathtt{N}_n}\sum_{k=1}^{\mathtt{N}_n} V_k\right).$$

The term in front of the average goes to 1 with probability one, and the the average goes almost surely to $p$ by the strong law of large numbers applied to the Bernoulli random variables $B_k$ (since $\mathtt{N}_n \to +\infty$ with probability one).

Finally, for the convergence in law, we exploit again the previous results, which can be gathered as :

$$\frac{\mathtt{N}_n - a_n^{(d)}}{b_n^{(d)}} \xrightarrow{\text{law}} \mathcal{W}^{(d)} \tag{B.30}$$

where

$$\mathcal{W}^{(1)} = \max_{0\le s\le 1} \mathcal{B}_s - \min_{0\le s\le 1} \mathcal{B}_s, \quad a_n^{(1)} = 0, \quad b_n^{(1)} = \sqrt{n},$$
$$\mathcal{W}^{(2)} = -2\pi^2 \mathcal{L}_{\mathcal{G}}, \quad a_n^{(2)} = \frac{\pi n}{\log n}, \quad b_n^{(2)} = \frac{n}{(\log n)^2},$$
$$\mathcal{W}^{(3)} = \mathcal{N}(0,1), \quad a_n^{(3)} = \kappa(3)n, \quad b_n^{(3)} = \sigma\sqrt{n \log n},$$

see (A.9), (A.17), and (A.22).

We use Lévy's convergence theorem to get convergence in law for $\mathtt{N}_n^{(p)}$ from the one for $\mathtt{N}_n$. More precisely, (B.28) yields

$$\mathtt{N}_n^{(p)} = p\ \mathtt{N}_n + \sum_{k=1}^{\mathtt{N}_n} (V_k - p).$$

Since $\mathbb{E}\left(\mathrm{e}^{\lambda(V_k-p)}\right) = (1-p)\,\mathrm{e}^{-\lambda p} + p\,\mathrm{e}^{\lambda(1-p)}$ ($\lambda \in \mathbb{C}$), and conditioning upon $\mathtt{N}_n$, we get

$$\begin{aligned}
\mathbb{E}\left(\mathrm{e}^{\mathrm{i}t(\mathtt{N}_n^{(p)} - pa_n^{(d)})/b_n^{(d)}}\right) &= \mathrm{e}^{-\mathrm{i}tpa_n^{(d)}/b_n^{(d)}}\ \mathbb{E}\left(\left((1-p)\,\mathrm{e}^{-\mathrm{i}tp/b_n^{(d)}} + p\,\mathrm{e}^{\mathrm{i}t(1-p)/b_n^{(d)}}\right)^{\mathtt{N}_n} \mathrm{e}^{\mathrm{i}tp\ \mathtt{N}_n/b_n^{(d)}}\right)\\
&= \mathrm{e}^{-\mathrm{i}tpa_n^{(d)}/b_n^{(d)}}\ \mathbb{E}\left(\mathrm{e}^{\left(\mathrm{i}tp/b_n^{(d)} + o(1)\right)\ \mathtt{N}_n}\right)\\
&= \mathbb{E}\left(\mathrm{e}^{\mathrm{i}tp(\mathtt{N}_n - a_n^{(d)})/b_n^{(d)}}\right) + o(1)\\
&\xrightarrow[n\to+\infty]{} \mathbb{E}\left(\mathrm{e}^{\mathrm{i}tp\mathcal{W}^{(d)}}\right)
\end{aligned}$$

using (B.30). By Lévy's convergence theorem, this gives

$$\frac{\mathtt{N}_n^{(p)} - p\,a_n^{(d)}}{b_n^{(d)}} \xrightarrow{\text{law}} p\,\mathcal{W}^{(d)}\,.$$

### C Derivation of the coefficient of variation

#### C.1 Reminder: Distribution of the consumption rate

The distribution of the number of prey consumed by a single predator after a time $t$ is given in App. B. Recall that the consumption rate (or functional response) after a time $t$ is defined as

$$F_t^{(d)} := \frac{R_t^{(d)}}{t}$$

which gives

$$F_t^{(d)} = \begin{cases} \frac{p}{\sqrt{\tau_e t}}\,\mathcal{W}_t^{(1)} + o_{\mathbb{P}}\left(\frac{1}{\sqrt{t}}\right) & \text{if } d=1 \\ \frac{\pi p}{\tau_e \ln t/\tau_e}\left(1 - \frac{1}{\ln t/\tau_e}\left(\gamma + \pi\frac{\tau_h}{\tau_e} + 2\pi\,\mathcal{W}_t^{(2)}\right)\right) + o_{\mathbb{P}}\left((\frac{1}{\ln t})^2\right) & \text{if } d=2 \\ \frac{\kappa p}{\tau_e + \kappa\tau_h} + p\sqrt{\frac{\ln t/\tau_e}{t\tau_e(1+\kappa\tau_h/\tau_e)^3}}\,\mathcal{W}_t^{(3)} + o_{\mathbb{P}}\left(\sqrt{\frac{\ln t}{t}}\right) & \text{if } d=3 \end{cases} \tag{C.1}$$

where $p$ is the probability that a site contains a prey, $\gamma \approx 0.577$ is Euler-Mascheroni constant, $\kappa \approx 0.659$ is the probability that a simple symmetric random walk never returns to the origin in $\mathbb{Z}^3$, and $\mathcal{W}_t^{(d)} =_{\text{law}} \mathcal{W}^{(d)}$ are different random variables depending on dimensionality (see App. B.2 for details): $\mathcal{W}^{(1)}$ is the difference between the max and min position of a 1-dimensional Brownian motion: $\mathcal{W}^{(1)} = \max_{s\leq 1} \mathcal{B}_s^{(1)} - \min_{s\leq 1} \mathcal{B}_s^{(1)}$; $W^{(2)} = \mathcal{L}_{\mathcal{G}}$ is the local time of self-intersection of a 2-dimensional Brownian motion; $\mathcal{W}^{(3)} = \mathcal{N}(0,\eta^2)$ is a centred Gaussian distribution with a constant variance $\eta^2$. Finally, $o_{\mathbb{P}}(\eta_t)$ is defined such that for any $\varepsilon > 0$, $\mathbb{P}(o_{\mathbb{P}}(\eta_t) \geq \varepsilon\eta_t) \to 0$ as $t \to \infty$.

The mean, variance and the coefficient of variation of the consumption rate $F_t$ are summarized and illustrated in the following table and figures:

Table 1: Summary of statistics for the distribution of the consumption rate.

| Dimension | $\mathbb{E}$xpectation | Varariance | Coefficient of variation |
|---|---|---|---|
| 1D | $2p\sqrt{\frac{2}{\pi}}\frac{1}{\sqrt{\tau_e t}}$ | $4p^2\frac{\ln 2 - 2/\pi}{t\tau_e}$ | $\sqrt{\frac{\pi}{2}\ln 2 - 1} \approx 0.298$ |
| 2D | $p\frac{\pi/\tau_e}{\ln t/\tau_e}\left(1 - \frac{\gamma+\tau_h\pi/\tau_e}{\ln t/\tau_e}\right)$ | $p^2\frac{2K\pi^2}{\tau_e^2(\ln t/\tau_e)^4}$ | $\frac{\sqrt{2K}}{\log\frac{t}{\tau_e} - \gamma - \pi\frac{\tau_h}{\tau_e}} \approx \frac{1.3}{\log t/\tau_e}$ |
| 3D | $p\frac{\kappa(3)}{\tau_e+\kappa(3)\tau_h}$ | $p^2\frac{27\kappa(3)^4}{2\pi^2}\frac{\ln t/\tau_e}{t\tau_e(1+\kappa(3)\tau_h/\tau_e)^3}$ | $\frac{0.8}{\sqrt{1+0.7\frac{\tau_h}{\tau_e}}}\sqrt{\frac{\log\frac{t}{\tau_e}}{\frac{t}{\tau_e}}}$ |

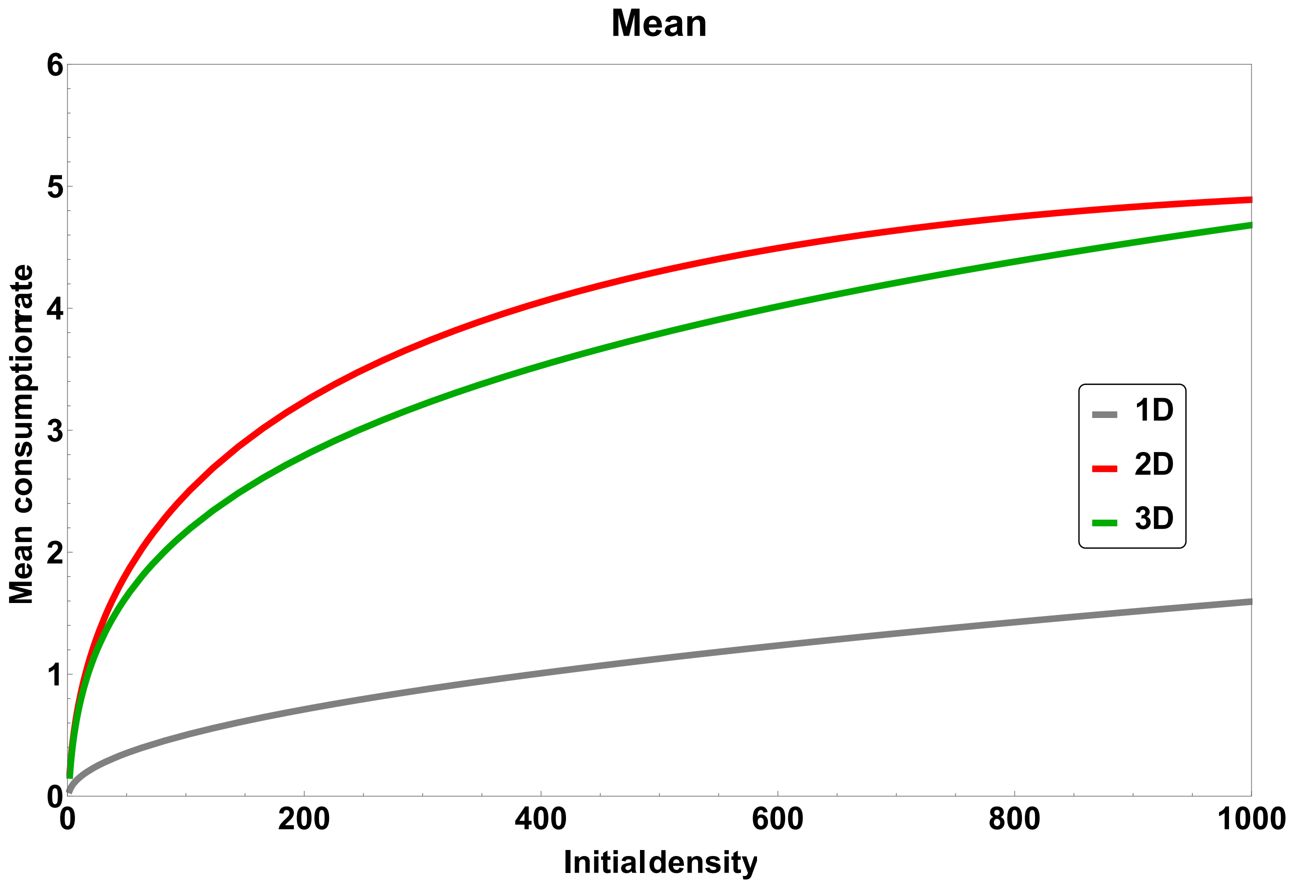


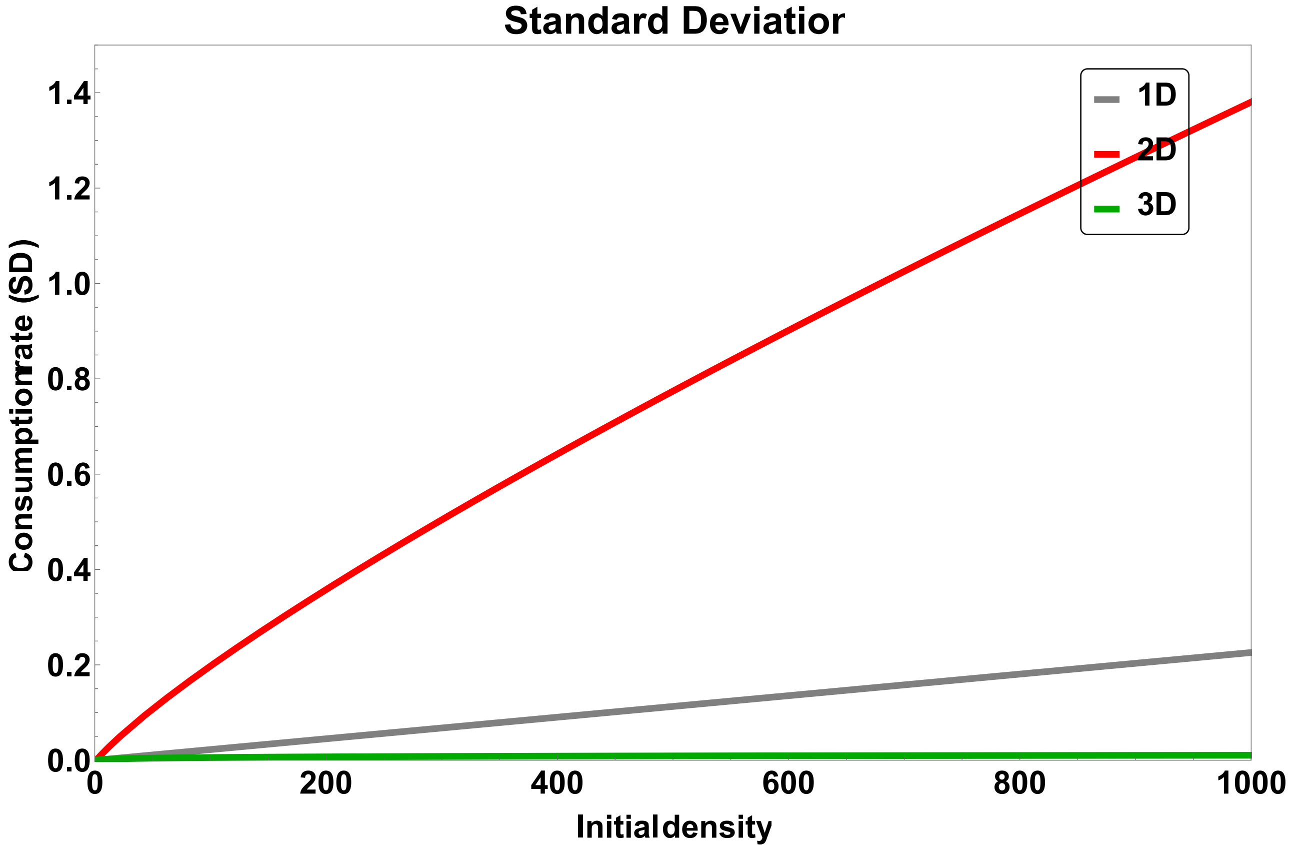


Figure C.1: Illustration of the mean and standard deviation of the consumption rate $F_t$ (parameters: $t = 10^3, \tau_e = 1, \tau_h = 0.045$).

#### C.2 Coefficient of variation of the consumption rate with between individuals variability

Suppose now that the consumption rate is a random variable which depends on two independent sources of random variations denoted $\omega_1$ and $\omega_2$, such that $F_t(\omega_1, \omega_2) := R_t(\omega_1, \omega_2)/t$. $\omega_1$ is the between-individual variability for searching and handling , *i.e.* parameters involved in searching $\tau_e(\omega_1)$ and handling $\tau_h(\omega_1)$ for a focal individual are randomly chosen in distributions depending on $\omega_1$. The parameters involved in the searching and handling times can have any explicit form (not necessarily Gaussian) but with finite mean and variance. $\omega_2$ is the variability due to the foraging stochastic process itself, *i.e.* the stochastic trajectory of the focal individual when foraging follows a random distribution $\mathcal{W}^{(d)}(\omega_2)$. The distribution $\mathcal{W}^{(d)}(\omega_2)$ followed by the trajectories directly emerges from the assumptions of the foraging process, here the simple random walk in dimension $d$.

As shown in (C.1), when the foraging time $t$ is large, the distribution of the consumption rate in a $d$-dimension space after a time $t$ for a focal individual takes the general form

$$F_t^{(d)}(\omega_1, \omega_2) = m_t^{(d)}(\omega_1) + s_t^{(d)}(\omega_1)\mathcal{W}_t^{(d)}(\omega_2) + o_{\mathbb{P}}\left(s_t^{(d)}(\omega_1)\right), \tag{C.2}$$

where $m_t^{(d)}(\omega_1)$ and $s_t^{(d)}(\omega_1)$ are functions which depend on time $t$, the dimension of space $d$, and the properties of individuals given by parameters $\tau_h(\omega_1)$ and $\tau_e(\omega_1)$. When these parameters are compactly supported in $(0, \infty)$, the proof of uniform integrability, namely Proposition B.4, can be directly extended, since all bounds are uniform with respect to parameters. We obtain that the mean and variance of the consumption rate satisfy, for $t \to \infty$ (thereafter notations $\omega_1$ and $\omega_2$ are dropped for the sake of simplicity):

$$\begin{aligned}
\mathbb{E}\left[F_t^{(d)}\right] &\sim \mathbb{E}\left[m_t^{(d)}\right] + \mathbb{E}\left[s_t^{(d)}\mathcal{W}^{(d)}\right], \\
\mathrm{Var}\left[F_t^{(d)}\right] &\sim \mathbb{E}\left[\left(m_t^{(d)} + s_t^{(d)}\mathcal{W}^{(d)}\right)^2\right] - \mathbb{E}\left[\left(m_t^{(d)} + s_t^{(d)}\mathcal{W}^{(d)}\right)\right]^2 \\
&\sim \mathbb{E}\left[\left(m_t^{(d)}\right)^2\right] - \mathbb{E}\left[\left(m_t^{(d)}\right)\right]^2 \\
&\quad + \mathbb{E}\left[\left(s_t^{(d)}\mathcal{W}^{(d)}\right)^2\right] - \mathbb{E}\left[\left(s_t^{(d)}\mathcal{W}^{(d)}\right)\right]^2 \\
&\quad + 2\left(\mathbb{E}\left[\left(m_t^{(d)}s_t^{(d)}\mathcal{W}^{(d)}\right)\right] - \mathbb{E}\left[m_t^{(d)}\right]\mathbb{E}\left[s_t^{(d)}\mathcal{W}^{(d)}\right]\right).
\end{aligned}$$

As the sources of variability $\omega_1$ and $\omega_2$ are independent, we have

$$\begin{aligned}
\mathbb{E}\left[s_t^{(d)}\mathcal{W}^{(d)}\right] &= \mathbb{E}\left[s_t^{(d)}\right]\mathbb{E}\left[\mathcal{W}^{(d)}\right], \\
\mathrm{Var}\left[s_t^{(d)}\mathcal{W}^{(d)}\right] &= \mathbb{E}\left[\left(s_t^{(d)}\right)^2\right]\mathrm{Var}\left[\mathcal{W}^{(d)}\right] + \mathrm{Var}\left[s_t^{(d)}\right]\mathbb{E}\left[\mathcal{W}^{(d)}\right]^2.
\end{aligned}$$

In addition, as we have $\mathbb{E}\left[m_t^{(1)}\right] = 0$ and $\mathbb{E}\left[\mathcal{W}^{(2)}\right] = \mathbb{E}\left[\mathcal{W}^{(3)}\right] = 0$ (Eqs. C.1), the expression of the variance of the consumption rate $F_t^{(d)}$ further simplifies to

$$\operatorname{Var}\left[F_t^{(d)}\right] \sim \mathbb{E}\left[\left(m_t^{(d)}\right)^2\right] - \mathbb{E}\left[\left(m_t^{(d)}\right)\right]^2 + \mathbb{E}\left[\left(s_t^{(d)}\mathcal{W}^{(d)}\right)^2\right] - \mathbb{E}\left[\left(s_t^{(d)}\mathcal{W}^{(d)}\right)\right]^2 \tag{C.3}$$

$$\sim \operatorname{Var}\left[m_t^{(d)}\right] + \operatorname{Var}\left[s_t^{(d)}\mathcal{W}^{(d)}\right]. \tag{C.4}$$

Finally, a general expression for the coefficient of variation of the consumption rate is

$$\operatorname{CV}\left[F_t^{(d)}\right] \sim \frac{\sqrt{\operatorname{Var}\left[m_t^{(d)}\right] + \operatorname{Var}\left[s_t^{(d)}\mathcal{W}^{(d)}\right]}}{\mathbb{E}\left[m_t^{(d)}\right] + \mathbb{E}\left[s_t^{(d)}\mathcal{W}^{(d)}\right]}$$

$$\sim \frac{\sqrt{\operatorname{Var}\left[m_t^{(d)}\right] + \mathbb{E}\left[\left(s_t^{(d)}\right)^2\right]\operatorname{Var}\left[\mathcal{W}^{(d)}\right] + \operatorname{Var}\left[s_t^{(d)}\right]\mathbb{E}\left[\mathcal{W}^{(d)}\right]^2}}{\mathbb{E}\left[m_t^{(d)}\right] + \mathbb{E}\left[s_t^{(d)}\right]\mathbb{E}\left[\mathcal{W}^{(d)}\right]}$$

which finally gives (Eqs. C.1)

$$\begin{cases} \operatorname{CV}\left[F_t^{(1)}\right] & \sim \frac{\sqrt{\mathbb{E}\left[\left(s_t^{(1)}\right)^2\right]\operatorname{Var}\left[\mathcal{W}^{(1)}\right] + \operatorname{Var}\left[s_t^{(1)}\right]\mathbb{E}\left[\mathcal{W}^{(1)}\right]^2}}{\mathbb{E}\left[s_t^{(1)}\right]\mathbb{E}\left[\mathcal{W}^{(1)}\right]} = \frac{1}{\mathbb{E}\left[s_t^{(1)}\right]}\sqrt{\operatorname{Var}\left[s_t^{(1)}\right] + \mathbb{E}\left[\left(s_t^{(1)}\right)^2\right]\frac{\operatorname{Var}\left[\mathcal{W}^{(1)}\right]}{\mathbb{E}\left[\mathcal{W}^{(1)}\right]^2}}, \\ \operatorname{CV}\left[F_t^{(2)}\right] & \sim \frac{\sqrt{\operatorname{Var}\left[m_t^{(2)}\right] + \mathbb{E}\left[\left(s_t^{(2)}\right)^2\right]\operatorname{Var}\left[\mathcal{W}^{(2)}\right]}}{\mathbb{E}\left[m_t^{(2)}\right]} = \frac{1}{\mathbb{E}\left[m_t^{(2)}\right]}\sqrt{\operatorname{Var}\left[m_t^{(2)}\right] + \mathbb{E}\left[\left(s_t^{(2)}\right)^2\right]\operatorname{Var}\left[\mathcal{W}^{(2)}\right]}, \\ \operatorname{CV}\left[F_t^{(3)}\right] & \sim \frac{\sqrt{\operatorname{Var}\left[m_t^{(3)}\right] + \mathbb{E}\left[\left(s_t^{(3)}\right)^2\right]\operatorname{Var}\left[\mathcal{W}^{(3)}\right]}}{\mathbb{E}\left[m_t^{(3)}\right]} = \frac{1}{\mathbb{E}\left[m_t^{(3)}\right]}\sqrt{\operatorname{Var}\left[m_t^{(3)}\right] + \mathbb{E}\left[\left(s_t^{(3)}\right)^2\right]\operatorname{Var}\left[\mathcal{W}^{(3)}\right]}. \end{cases}$$

Roughly speaking, the coefficient of variation of the consumption rate $F_t^{(d)}$ is a function of two variance components such as: $c_1 \operatorname{Var}[\omega_1] + c_2 \operatorname{Var}[\omega_2] + c_3 \operatorname{Var}[\omega_1]\operatorname{Var}[\omega_2]$, where $c_1, c_2, c_3$ are constants. This shows that if the between-individuals variability has a lower order than the variability of the foraging process itself ($\operatorname{Var}[\omega_1] \ll \operatorname{Var}[\omega_2]$), then the coefficient of variation of the consumption rate $\operatorname{CV}[F_t]$ should be of the same order than the coefficient of variation of the foraging process, *i.e.* of order 1 in 1D, $1/\ln t$ in 2D, or $1/\sqrt{t}$ in 3D. In other words, if the main driver of the variation of the consumption rate is the foraging process itself, then the coefficients of variation estimated from data should be at most of order 1 (or more precisely it should fall within the range given in Tab. 1). If its order of magnitude is larger than 1 (or more precisely if it falls outside the range given in Tab. 1), then we could conclude that the main driver of the variation of the consumption rate is the between-individual variability (*i.e.* $\operatorname{Var}[\omega_1] \gg \operatorname{Var}[\omega_2]$).

Two other alternatives are possible, yet less plausible. First, the coefficient of variation estimated from data could be of order lower than 1, which would mean that both the between-individual variability and the variability due to the foraging process itself are of order lower than 1. Second, the coefficient of variation estimated from data could be of order 1 and at the same time the between-individual variability be of order larger than the variability from the foraging process. If so, it would remain to explain how it is possible that the between-individual variability is such that the coefficient of variation falls exactly in this order of magnitude. Those two alternatives would seem even less plausible when the estimated coefficients of variation cover a large number of species and experimental and ecological contexts, as in our case and the datasets we compiled.

Therefore, we finally make two global predictions: either the coefficients of variation estimated from data are at most of order 1, or they are of order larger than 1. We would then conclude that the main driver of the variation of the consumption rate is the foraging process in the former case, or that it is external sources of variations, such as the between-individual variability, in the latter.

### D Stochastic simulations

#### D.1 Standard setup

We first evaluated the accuracy and the convergence rate of the approximation of $CV_t^{(d)}$, the coefficient of variation of the consumption rate, defined as the number of prey per unit of time consumed by a predator foraging in a $d$-dimensional space during a total foraging duration $t$.

To achieve this, we conducted numerical stochastic simulations of a consumer randomly and symmetrically walking on a $d$-dimensional homogeneous square lattice where each node initially contains a prey (Fig. D.1). The consumer moves from one node to any of the four other adjacent nodes at constant speed $v$. When a node with a prey is encountered, the consumer feeds on the prey, which takes a constant handling time $\tau_h$, and the node is depleted (we assume there is no regeneration of the prey). If the node was already visited, the consumer randomly moves to any of the adjacent node without spending handling time on the currently visited node.

Denoting $x$ the initial density of prey in the lattice, and $L$ an arbitrary length unit, the distance between two prey in the lattice is

$$y^{(d)}(x) = \frac{L}{x^{1/d} - 1}. \tag{D.1}$$

The time to cover this distance is then given by $\tau_e^{(d)}(x) = y^{(d)}(x)/v$. A simulation run (i.e. one trajectory or realization of the stochastic process) is stopped when the total foraging duration $t$ is reached, time at which the total number of prey consumed $R_t^{(d)}(x)$ is counted, and the consumption rate $F_t^{(d)}(x) = R_t^{(d)}(x)/t$ is recorded. $10^4$ realizations per initial density $x$ were run with the following parameters values: $t = 10^7$, $v = 1$, $\tau_h = 0.1$, $L = 1000$.

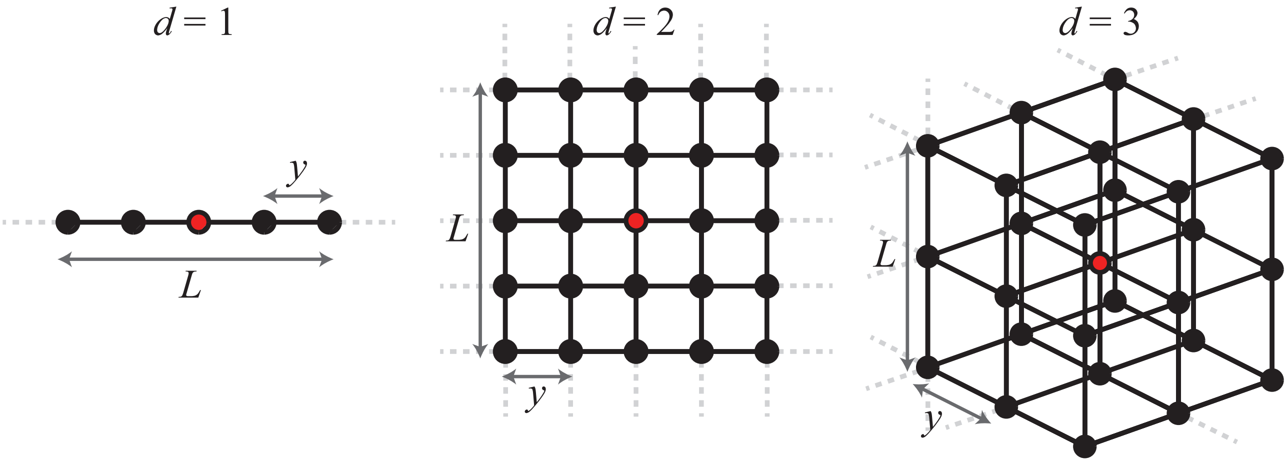


Figure D.1: Illustration of the lattice for $d = \{1, 2, 3\}$. The red dot denotes the starting position of the walker. $L$ is an arbitrary length unit; $y$ is the distance between two nodes; $d$ is the dimension of space.

#### D.2 Robustness

We evaluated, under various ecological contexts, the robustness of the analytical approximations of $CV_t^{(d)}$, the coefficient of variations of the consumption rate. We ran numerical stochastic simulations of modified versions of the standard algorithm described in the previous section. Specifically, we ran simulations with one the four following additional mechanisms (detailed below): Walk with stochastic length jumps, walk with drift, self-avoiding walk, and walk in an initially heterogeneous and random repartition of prey on the lattice.

**Jumps with random length.** The consumer moves in the lattice following a symmetrical random walk with jumps of length following a truncated power-law distribution $P(k) \sim k^{-\theta}$ with $k \in [1, k_{\max}]$, where $k_{\max}$ is the maximum step-size. Simulations were run for the following parameter values: $\theta \in \{2.0, 2.5, 3.0, 3.5\}$, $k_{\max} = 10$. Other parameters were the same than in the standard setup (see previous section).

**Walk with drift.** The consumer moves on the lattice in a preferred direction, denoted $\rightarrow$. The drift parameter $\mu \in [0, 1]$ is defined such that, each step, the consumer has the probability

$$\mathbb{P}(\text{jump} =' \rightarrow') = \frac{1 + (2d - 1)\mu}{2d}$$

to move in this preferred direction, and a probability

$$\mathbb{P}(\text{jump} \neq' \rightarrow') = \frac{1 - \mu}{2d}$$

to move in any of the $2d - 1$ remaining directions: $\leftarrow$ in 1D, $\leftarrow$, $\downarrow$ or $\uparrow$ in 2D, $\leftarrow$, $\downarrow$ or $\uparrow$, $+1\,level$ or $-1\,level$ in 3D. If $\mu = 0$ the movement is symmetrical and resumes to the standard random walk. If $\mu = 1$, the movement is unidirectional. Simulations were run with following parameter values, corresponding to an increasing biased random walk $\mu \in \{0.05, 0.1, 0.25, 0.5\}$.

**Self-avoiding walk with a short term memory.** The consumer had a short-term memory avoiding the node it just left.

**Walk on a lattice with a random initial repartition of prey.** Each node of the lattice initially contains a prey with probability $p$. In practice, each time the consumer visited an unvisited node, a Bernoulli trial with probability $p$ were drawn. If $p = 1$, the walk resumes to a typical random walk on a uniform graph (as detailed in the previous section), if $p = 0$ the lattice is initially empty. By default simulations were run for $p = 1/2$, unless indicated.

### E Data

N.B.: All data, associated papers (when available) and R scripts are available on the following server: github.

#### E.1 Dataset 1: FoRAGE database

The first dataset was obtained from the database FoRAGE [15], dowloaded from https://knb.ecoinformatics.org/ in january 2021 (version 12.18.18). The database contains the predator and prey species names, the literature source, the initial prey density (either per $m^2$ or $m^3$), the consumption rate (the number of prey eaten per day per predator), the sample size per treatment (i.e. the initial density). The database FoRAGE was partly built by automatically extracting data from papers' figures, and partly populated directly from tables (see [15] for details). We discarded all data where at least one of these information was missing.

There were two categories of available data that were differently used for the calculation of the coefficient of variation estimations: i) raw data or ii) summary statistics. In the case of i) raw data: because there could have overlaps between dots in figures (i.e. when different predators fed on the same number of prey), the number of data could be lower than the sample size announced in the initial paper, in particular when the initial density of prey was low. We thus curated the database by removing the data with too large uncertainty: we only kept data where the sample size was $\geq 8$ and where the number of available measured consumption rate was at least 80% of the sample size (i.e. at least 8 data points if the sample size was 10, for example). In addition, when a single dot was detected from the figure for a given experimental treatment, the FoRAGE's authors filled the database with a single value for as many independent replicates as supposed from the methods described in the original paper (for example, if the sample size was $n = 10$ but a single dot in a figure was observed, the database was populated with 10 rows, one for each replicate, with the same number of eaten prey). As a consequence, there was no variation in the consumption rate in this case but with no certainty about whether it was due to the limit of data collection itself or because there was indeed no variation. We thus decided to exclude these data when there was no variation. The coefficients of variation were estimated by calculating the mean and the standard deviation within initial density directly from raw data. This resulted in 142 estimated coefficients of variation (calculated on 1266 from 15108 initial data after curation), for 19 predator and 17 prey species.

In the case of ii) summary statistics: we only kept data where the mean, the standard error and the sample size were available, where the sample size was $\geq 8$ (to ensure minimal quality for summary statistics estimations and be consistent with case i) raw data), and where the standard error was $> 0$ (because there was uncertainty whether it was true or false zero, and because a standard error equal to 0 was mostly encountered in low densities treatments suggesting that all prey were eaten in all replicates). We calculated the coefficients of variation directly from the mean, standard error and sample size, within an initial density, by applying the formula of the definition of the standard error: $\sigma = \sqrt{n}\ se$ (where $n$ is the sample size, $\sigma$ is the standard deviation, and $se$ is the standard error;

$n$ and $se$ are given in the database). This resulted in 3039 estimated coefficients of variation aftr curation (from 6797 estimated mean and $se$ consumption rates in total in the database), for 126 predator and 104 prey species.

Overall the coefficients of variation calculated from data compiled in the FoRAGE database may contain the following sources of variations: i) between individuals; ii) within individuals (if different or identical individuals were used for a given initial density or for different initial densities, information not available from the database); iii) between environments, as it was not known to what extent the environmental conditions of the experiment were controlled; iv) experimental and observations errors; v) errors due to the FoRAGE database build itself (in particular due to the automatic image analysis and data collection); vi) from the foraging process itself.

Fig. E.1 shows the median and quartiles of all the coefficients of variation calculated from the FoRAGE database. The median is 0.2905. 73.2% of the coefficients of variation lie between 0.1 and 1, and 95.5% lie between 0.01 and 10.

#### E.2 Dataset 2: Raw data, potentially entangled between- and within- individuals variations

The datasets were obtained from

- The Dryad deposit (search in january 2021 with keywords: "foraging", "intake", "functional response", "visitation rate", 15 collected datasets).
- The database by [16] available at `github.com/stoufferlab/general-functional-responses` (25 datasets). From this database, we extracted and used only datasets where the number of predators was fixed to one, where there was several measurements at a given density, and a single type of prey; we excluded any dataset where inconsistencies were detected or with uncertainties; we included datasets where the mean, the sample size and the standard errors were available, from which we calculated the coefficients of variation.
- Data from direct authors sharing. We directly contacted authors for data sharing (1 dataset from [17].

Overall we computed 602 coefficients of variation from 41 different datasets (from which 20 coefficients of variation were equal to 0, all in the case of low densities: suggesting that all prey were consumed by all predators, resulting in a standard deviation equal to 0; As the minimum non-zero value of the coefficient of variation was $\geq 0.029$, we discarded zero values). The coefficients of variation calculated from these datasets potentially contains the following sources of variations: i) between individuals; ii) within individuals (if different or identical individuals were used for a given initial density or for different initial densities, information not available from the database by [16], or with uncertainty from datasets downloaded from Dryad, or the dataset shared by the author ); iii) between environments, as it was not known to what extent the environmental conditions of the experiment were controlled; iv) experimental and observations errors; v) the foraging process itself.

Fig. E.1 shows the median and quartiles of all the coefficients of variation calculated from these raw datasets. The median is 0.408, 87.8% of the coefficients of variation lie between 0.1 and 1, and 100% lie between 0.01 and 10.

List of references: Data obtained directly from the authors: [17]. Data obtained from Dryad deposit: [18, 19, 20, 21, 22, 23, 24, 25, 26, 27, 28, 29, 30, 31, 32]. Data from the database: [16].

##### E.3 Dataset 3: Raw data, accounting for within individuals variations

We collected three datasets directly from the authors where consumption rates were measured several times for the same individuals. One dataset were obtained in natural conditions [33], in semi-natural conditions [34], or in controlled conditions [35]. We directly calculated coefficients of variation within individuals from the raw data shared by [33] and [35]. The coefficients of variation were calculated by the authors themselves and shared with us in the case of the Imperial shag [34]. It resulted in 229 estimations of within-individuals coefficients of variation. In natural and semi-natural conditions, the sources of possible variations were: i) within individuals; ii) the environment (including uncontrolled prey densities); iii) experimental errors, iv) the foraging process itself. In the controlled conditions, the sources of possible variations were: i) within individuals; ii) experimental errors; iii) the foraging process itself. In the case of the crayfish experiments [35], we could calculate the coefficients of variation within individuals within and across densities as the same individuals were used several times for a given density, and for different densities.

Fig. E.1 shows the median and quartiles of all the coefficients of variation calculated from these datasets. The median is 0.273, 85.1% of the coefficients of variation lie between 0.1 and 1, and 99.1% lie between 0.01 and 10.

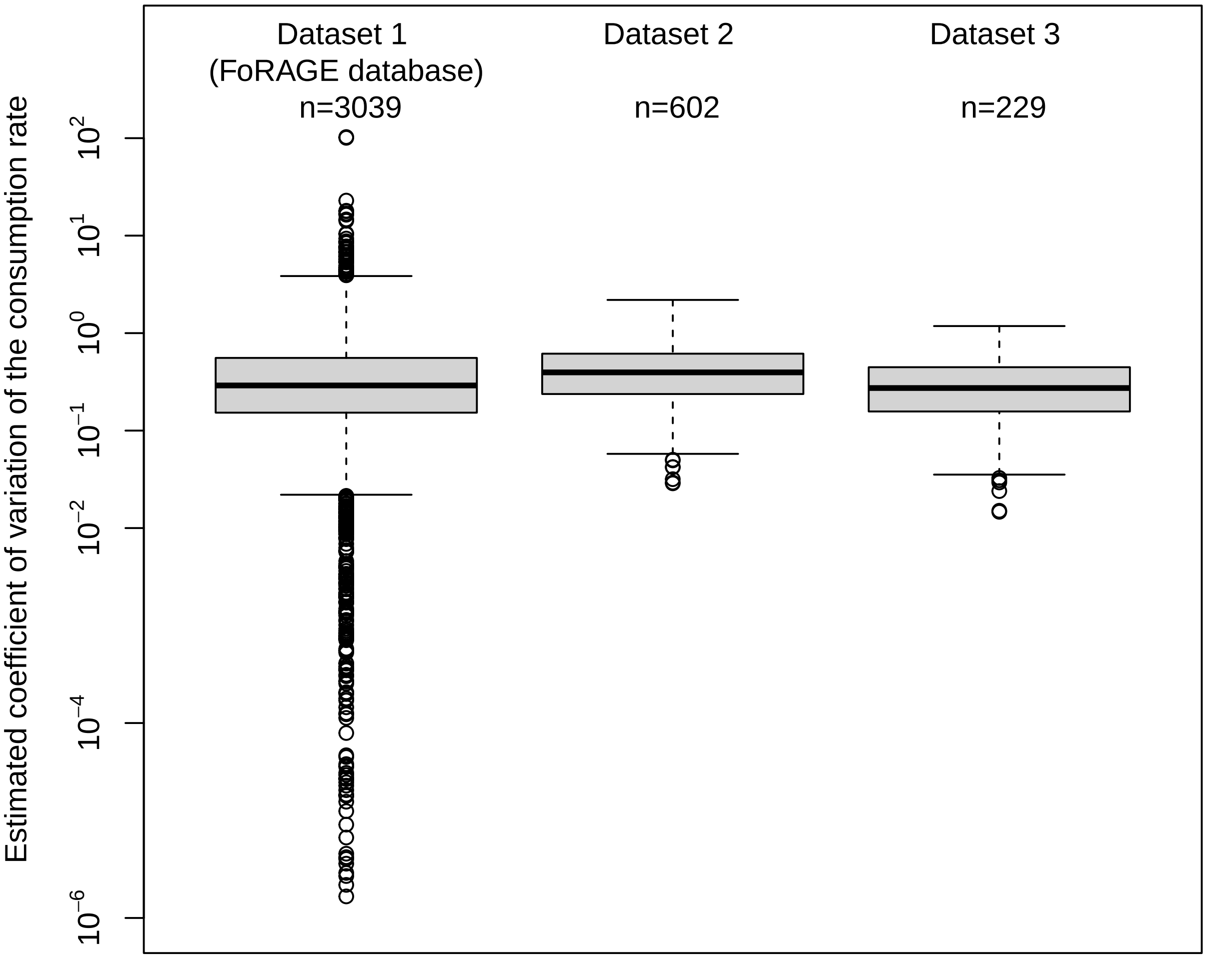


Figure E.1: Boxplot of the coefficients of variation of the consumption rate of prey per single predator calculated from different datasets (see text for details).

#### References

[1] Frank Spitzer. Principles of random walk, volume 34. Springer Science & Business Media, 2001.

[7] Enrico Masina. Useful review on the exponential-integral special function. arXiv preprint arXiv:1907.12373, 2019.

[8] J-F Le Gall. Propriétés d'intersection des marches aléatoires. Communications in mathematical physics, 104(3):471–507, 1986.

[9] Jean-François Le Gall. Sur le temps local d'intersection du mouvement brownien plan et la méthode de renormalisation de varadhan. In Séminaire de Probabilités XIX 1983/84, pages 314–331. Springer, 1985.

[10] Richard F Bass, Xia Chen, and Jay Rosen. Moderate deviations for the range of planar random walks. American Mathematical Soc., 2009.

[11] Patrick Billingsley. Convergence of probability measures. John Wiley & Sons, 2013.

[12] Naresh C Jain and William E Pruitt. The range of transient random walk. Journal d'Analyse Mathématique, 24(1):369–393, 1971.

[13] Gregory F Lawler and Vlada Limic. Random walk: a modern introduction, volume 123. Cambridge University Press, 2010.
